# Acclimation to high daily thermal amplitude converts a defense response regulator into susceptibility factor

**DOI:** 10.1101/2024.08.22.609129

**Authors:** Marie Didelon, Justine Sucher, Pedro Carvalho-Silva, Matilda Zaffuto, Adelin Barbacci, Sylvain Raffaele

**Author notes:** MAS Seeds, 1 Rte de Saint-Sever, 40280 Haut-Mauco, France.

## Abstract

Acclimation enables plants to adapt to immediate environmental fluctuations, supporting biodiversity and ecosystem services. However, global changes are altering conditions for plant disease outbreaks, increasing the risk of infections by pathogenic fungi and oomycetes, and often undermining plant immune responses. Understanding the molecular basis of plant acclimation is crucial for predicting climate change impacts on ecosystems and improving crop resilience. Here, we investigated how *Arabidopsis thaliana* quantitative immune responses acclimates to daily temperature fluctuations. We analyzed responses to the fungal pathogen *Sclerotinia sclerotiorum* following three acclimation regimes that reflect the distribution areas of both species. Mediterranean acclimation, characterized by broad diurnal temperature amplitudes, resulted in a loss of disease resistance in three natural *A. thaliana* accessions. Global gene expression analyses revealed that acclimation altered nearly half of the pathogen-responsive genes, many of which were down-regulated by inoculation and associated with disease susceptibility. Phenotypic analysis of *A. thaliana* mutants identified novel components of quantitative disease resistance following temperate acclimation. Several of these mutants were however more resistant than wild type following Mediterranean acclimation. Notably, mutant lines in the NAC42-like transcription factor did not show a loss of resistance under Mediterranean acclimation. This resistance was linked to an acclimation-mediated switch in the repertoire of NAC42-like targets differentially regulated by inoculation. These findings reveal the rewiring of immune gene regulatory networks by acclimation and suggest new strategies to maintain plant immune function in a warming climate.

## INTRODUCTION

The ability of species to cope with rising temperatures is a crucial factor influencing range shifts and local extinctions, as their distribution and range boundaries closely align with temperature gradients. Evidence shows that plant species adapt to local environmental conditions through genetic variation (Fournier-Level et al., 2011b; Katz et al., 2021; Clauw et al., 2022) but they also exhibit phenotypic plasticity allowing individual plants to adjust rapidly their physiology to environmental variations (Valladares et al., 2014; Brancalion et al., 2018). The short-term, reversible process that allows plants to cope with immediate environmental fluctuations is often referred to as acclimation (Kleine et al., 2021). Plant acclimation help maintain the balance of natural systems, supporting biodiversity and the services that ecosystems provide, such as carbon sequestration and water regulation. With climate change modifying the distribution area of plants (Sloat et al., 2020) and causing more frequent and severe weather events (Newman and Noy, 2023), knowledge of how plants acclimate can inform strategies to manage ecosystems and agriculture. In this context, crops that can acclimate effectively are more likely to maintain high yields despite stressors. A better understanding of the genetic underpinnings of plant acclimation is therefore crucial for predicting the impact of climate change on ecosystems and for improving crop resilience.

Acclimation distinguishes from adaptation for involving changes to the expression of the genome instead of heritable changes to genome sequences (Kleine et al., 2021). Epigenetic and transcriptional regulation mechanisms mediating somatic stress memory are important players in plant acclimation (Charng et al., 2023; Zuo et al., 2023; Hadj-Amor et al., 2024). Cold acclimation, by which decreasing temperatures enhance freezing tolerance in plants involves alterations in membrane composition, the production of cryoprotective polypeptides and solutes, the activation of cold-responsive (COR) genes regulated by C-repeat binding transcription factors (CBFs/DREB1) (Liu et al., 2019). The accumulation of heat shock proteins (HSPs) regulated by heat shock transcription factors (HSFs) and histone 3 K4 methylation play a key role in heat acclimation (Kappel et al., 2023; Nishad and Nandi, 2021). Besides transcription factors and epigenetic marks, the hormone abscisic acid (ABA) is a central mediator of the accumulation of LEA-like protective proteins, stomatal closure and downregulation of photosynthesis under drought acclimation (Sadhukhan et al., 2022). Despite recent efforts, the interplay between regulatory mechanisms, molecular and phenotypic responses to plant acclimation is elusive.

With changes to the climate, not only the distribution range of plants changes, but also that of their enemies. Suitable conditions for plant disease outbreaks are expected to shift in time and space leading to a global poleward movement of plant pathogen geographic niches (Bebber et al., 2013) and an increased risk of infection by pathogenic fungi and oomycetes (Chaloner et al., 2021). Fungi, especially generalists with a broad range of plant hosts, are the most widespread and most rapidly spreading pathogens, so that if current rates persist, several major food producing countries would have fully saturated pathogen distributions by 2050 (Bebber et al., 2014). A paradigmatic example of such broad host range pathogen is the white and stem mold fungus *Sclerotinia sclerotiorum*, which infects hundreds of plant species and causes significant losses to vegetable and oil crops worldwide (Navaud et al., 2018; Peltier et al., 2012; Cohen, 2023). Although climate change may alter the overlap between crops cultivation area and *S. sclerotiorum* distribution range (Mehrabi et al., 2019), pathogen strains adapted to warm temperatures have been reported (Uloth et al., 2015) and extreme temperature may promote fungal development (Lane et al., 2019; Shahoveisi et al., 2022), raising concern about *Sclerotinia* disease incidence in the future (Singh et al., 2023).

Plant respond to *S. sclerotiorum* by activating quantitative disease resistance (QDR), an immune response involving multiple genes of weak to moderate phenotypic effect (Roux et al., 2014; Sucher et al., 2020). Molecular players involved in QDR against *S. sclerotiorum* include immune receptors, reactive oxygen species, phytohormones such as ABA, jasmonic acid and ethylene, transcription factors and phytoalexins (Perchepied et al., 2010; Mbengue et al., 2016; Derbyshire and Raffaele, 2023). Several of these determinants contribute to multiple biological processes such as plant development and response to the abiotic environment (Corwin et al., 2016; Badet et al., 2019; Léger et al., 2022). The genetic architecture of QDR suggests that the expression of many genes involved in QDR could be modulated by environmental conditions (Hadj-Amor et al., 2024), and that climate change may alter plant QDR response to *S. sclerotiorum* at the phenotypic and molecular level. Temperature increase notably is known to frequently impair plant immune responses, including QDR (Desaint et al., 2021; Aoun et al., 2017).

Analyses of plant immune responses under abiotic constraints generally focus on pathogen inoculation under prolonged and stable abiotic conditions. In *A. thaliana*, immunity against the bacterial pathogen *Pseudomonas syringae* pv. *tomato* at elevated temperature can be restored by the constitutive expression of *CBP60g*, a major transcriptional regulator of plant immunity genes and salicylic acid (SA) defense hormone production, downregulated by temperature (Kim et al., 2022). This finding indicates that engineering plant transcriptional circuits can mitigate the negative effect of climate change on some plant immune responses. Whether this strategy would restore resistance against necrotrophic pathogens such as *S. sclerotiorum*, only weakly sensitive to SA-mediated defense, remains to be determined. Another promising target is the disordered protein TWA1, a temperature sensor proposed to orchestrate acclimation by integrating temperature with ABA and JA signaling (Bohn et al., 2024), which play important roles in plant defense against necrotrophs. When applied sequentially, prior abiotic signals may alter the transcriptional and metabolic response to a subsequent pathogen inoculation (Coolen et al., 2016; Garcia-Molina et al., 2020; Garcia-Molina and Pastor, 2024). In addition to mean temperature increase, climate change drives an expansion of diurnal temperature range (Zhong et al., 2023). Daily fluctuations of the environment may alter plant metabolism, growth and flowering (Burghardt et al., 2016; Deng et al., 2021; Matsubara, 2018) as well as gene regulation and invasive growth of fungal pathogens (Jallet et al., 2020; Bernard et al., 2022). Yet, how plant immunity acclimates to daily temperature fluctuations remains largely unexplored.

To fill this gap, we analyzed *A. thaliana* immune responses upon *S. sclerotiorum* inoculation following three acclimation regimes representing the distribution area of these two species. Mediterranean acclimation, characterized by a broad diurnal temperature amplitude, caused a loss of disease resistance in the three natural accessions we tested. Using global gene expression analyses, we show that acclimation alters the expression of nearly a half of pathogen-responsive genes, many of which are down-regulated by inoculation and associated with disease susceptibility. The phenotypic analysis of *A. thaliana* mutants identified novel components of QDR following temperate acclimation. Several of these mutants were however more resistant than wild type following Mediterranean acclimation. In particular, contrary to wild type, two mutant lines in the NAC42-like transcription factor showed no loss of resistance upon Mediterranean acclimation. These phenotypes associated with a switch in the repertoire of NAC42-like targets differentially regulated by inoculation according to acclimation. These findings reveal the rewiring of immune gene regulatory networks by acclimation and open new perspectives to safeguard the functioning of the plant immune system in a warming climate.

## RESULTS

### Mediterranean-like acclimation impairs the resistance of several *A. thaliana* accessions to *S. sclerotiorum*

To determine the effect of acclimation on *A. thaliana* quantitative disease resistance (QDR), we analyzed phenotypic variation of three *A. thaliana* accessions after growth in three simulated climates. We selected accessions Col-0, Rld-2 and Shahdara (Sha) as representatives of genetic and geographical diversity of *A. thaliana* species. For acclimation, plants were grown under day length, day and night temperatures corresponding to the 30-year average for the month of April in areas with a temperate (Cfa), continental (Dfb) and Mediterranean (Csa) climates (**Fig. 1A, Fig. S1**). These correspond to climates in the distribution range of *A. thaliana* with major projected area variation by the end of this century (Alonso-Blanco et al., 2016; Peel et al., 2007; Cui et al., 2021). Plants were grown for 35 days under temperate and Mediterranean climate, corresponding to 13,405 and 13,930 °C.days, and for 70 days under continental climate corresponding to 11,480 °C.days before inoculation with *S. sclerotiorum* under infection-conducive conditions. Plant susceptibility was assessed using time-resolved automated phenotyping (Barbacci et al., 2020). After temperate acclimation, all accessions appeared similarly susceptible with only a slightly lower susceptibility (-9% average) for Rld-2 and a slightly higher susceptibility for Sha (+18% average) compared to Col-0 (**Fig. 1B, Table S1**). These phenotypes were not significantly altered upon continental acclimation. Mediterranean acclimation rendered all accession significantly more susceptible, with an average increase by 29% for Sha, 37% for Col-0 and 86% for Rld-2 as compared to temperate acclimation (**Fig 1B, C**). These results show that both genotype and acclimation affect the susceptibility of *A. thaliana* to *S. sclerotiorum* and that, regardless of genotype, Mediterranean acclimation caused the most significant loss of resistance.

**Fig 1.**
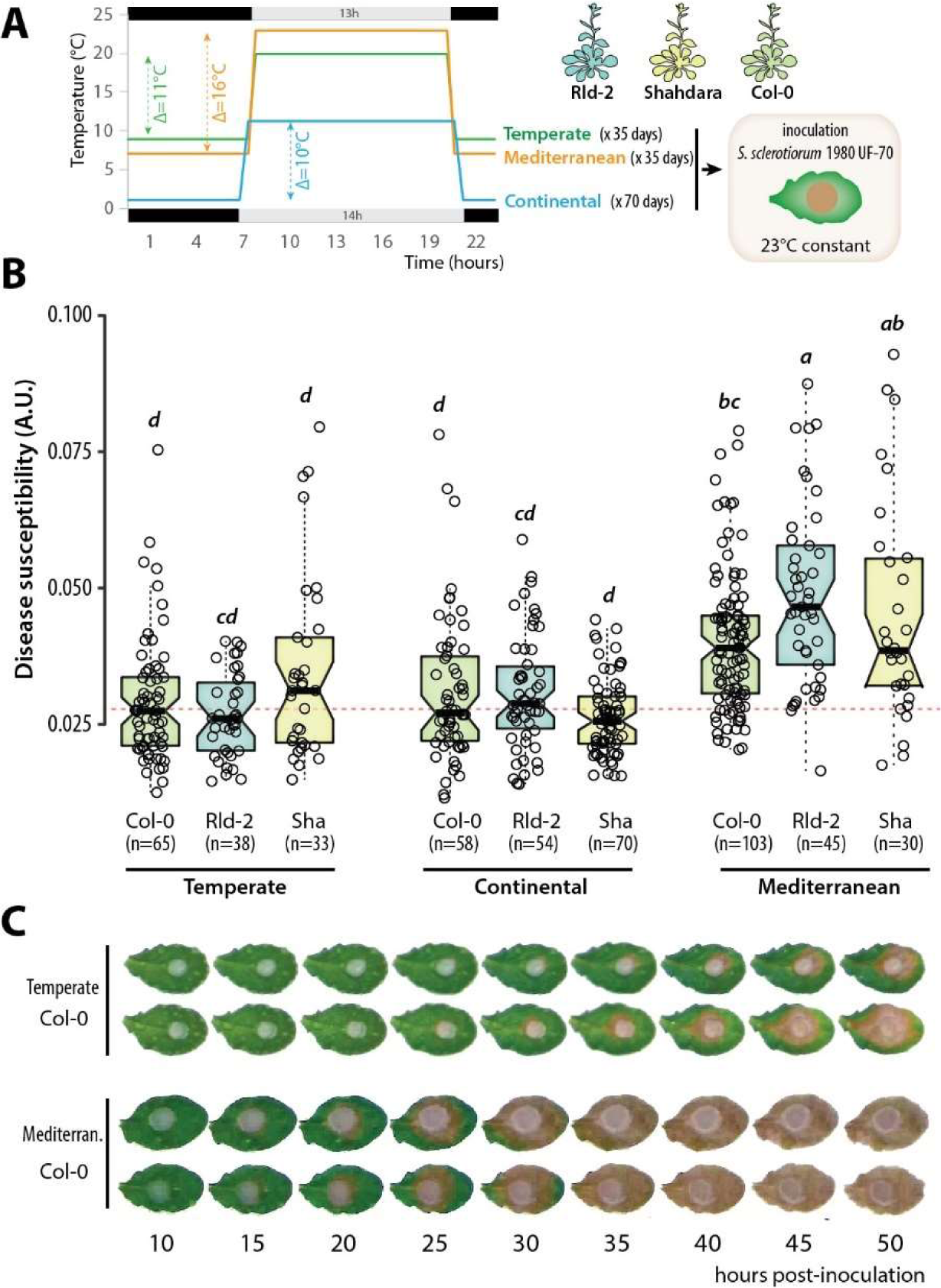
Effect of three distinct pre-infection climate conditions (acclimation) on *A. thaliana* quantitative disease resistance to *S. sclerotiorum*. (A) Experimental design showing acclimation and inoculation phases. Daylength, day and night temperatures typical of temperate, continental and Mediterranean climate conditions define the three acclimation conditions used in this work. (B) Susceptibility phenotype in response to *S. sclerotiorum* infection as a function of acclimation and genotype. Each experiment was repeated at least 3 times and the significance of the results was assessed by an ANOVA followed by a Tukey HSD test, with significance groups labelled by letters. Boxplots show first and third quartiles (box), median (thick line), and the most dispersed values within 1.5 times the interquartile range (whiskers). (C) Representative symptoms of Col-0 plants between 10 and 50 hours post-inoculation by *S. sclerotiorum* on leaves harvested on plants acclimated in temperate and Mediterranean (Mediterran.) conditions.

### Acclimation primarily alters the expression of infection-downregulated genes

To study the molecular bases of quantitative disease resistance acclimation, we performed a global transcriptome analysis of *A. thaliana* accessions Col-0, Rld-2 and Sha grown in temperate, continental and Mediterranean climates, followed or not by *S. sclerotiorum* inoculation. To identify genes responsive to infection we performed a differential expression analysis using non-inoculated plants as reference in each of nine conditions (three climate priming, times three plant genotypes). We found 17,137 nuclear-encoded genes with sufficient coverage (**Table S2**), and identified 9,580 differentially expressed <u>g</u>enes (DEGs) upon inoculation at |Log2 Fold Change|≥2 and Bonferroni-adjusted p-val<0.0001 (**Fig 2A, Fig S2, Table S3**). The number of upregulated genes ranged from 1,744 (Rld-2 Mediterranean acclimation) to 3,084 (Rld-2 continental acclimation), the number of downregulated genes ranged from 442 (Rld-2 Mediterranean acclimation) to 3,387 (Sha temperate acclimation). The three accessions showed a reduced number of DEGs when acclimated in conditions very divergent to the climate at their area of origin. Downregulated genes showed a relatively high degree of specificity with 1,212 genes (23%) unique to one genotype-acclimation pair and only 89 (1.7%) genes differential in all nine genotype-acclimation conditions tested (**Fig 2B**). Upregulated genes showed higher robustness with 1,105 genes (26.8%) differential in all nine genotype-acclimation conditions.

**Fig 2.**
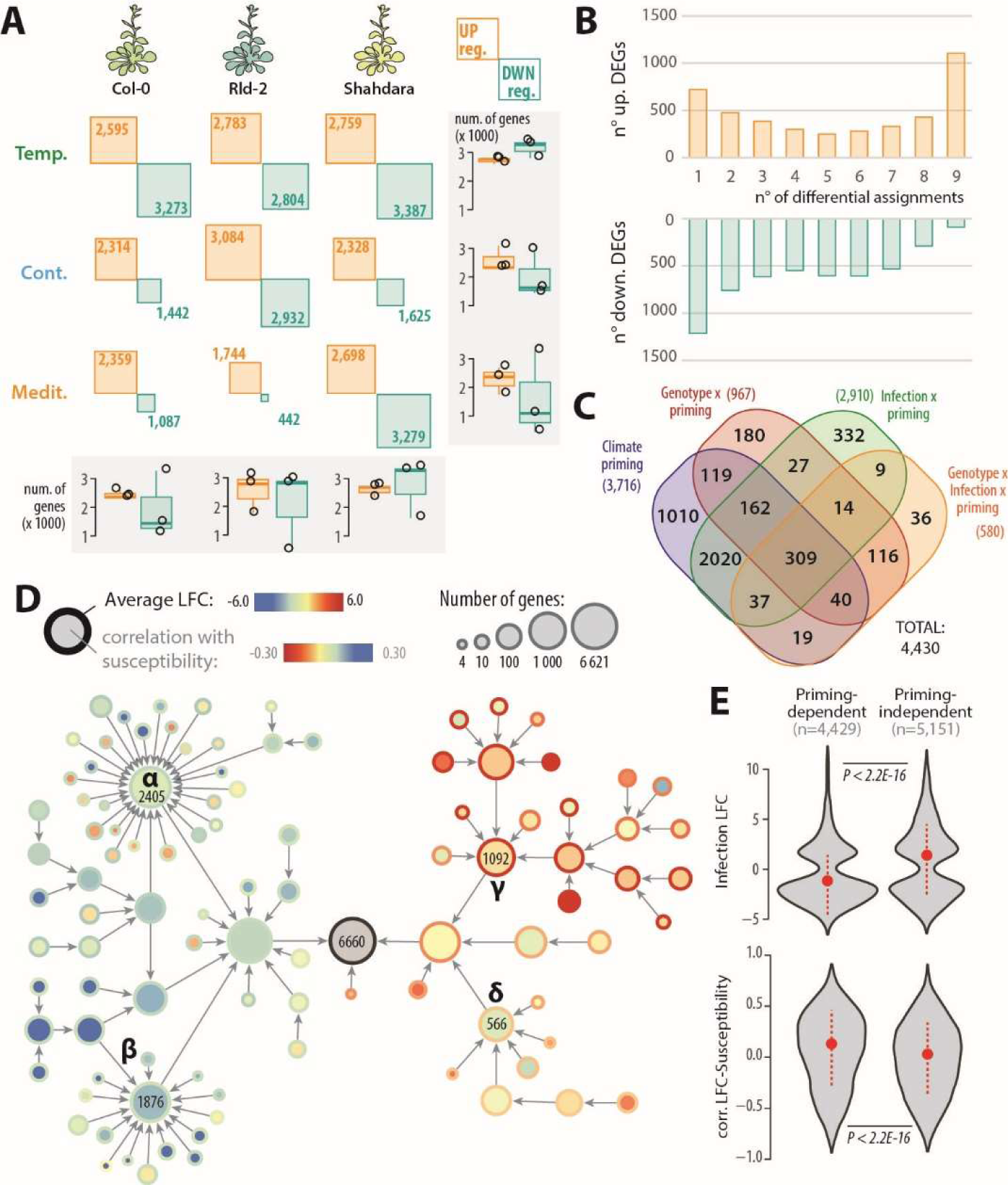
Global gene expression profiling of *A. thaliana* plants inoculated by the fungal pathogen *S. sclerotiorum* following temperate, continental and Mediterranean acclimation. (A) Number of differentially expressed genes (DEGs) upregulated (yellow) and down-regulated (blue) 48 hours post inoculation by *S. sclerotiorum* in each of three plant genotypes and three acclimation conditions. Box plots show the distribution of DEG number per genotype (columns) and acclimation (rows), first and third quartiles (box), median (thick line), and the most dispersed values within 1.5 times the interquartile range (whiskers) are shown. (B) Number of up- and down-regulated DEGs according to the number of differential assignments, out of 9 tested conditions. (C) Distribution of acclimation-dependent DEGs identified by ANOVA according to the factors explaining gene expression variance. (D) A hierarchical network of genes mis-regulated by *S. sclerotiorum* infection identified through differential and variance analyses. Nodes represent gene communities sized according to the number of DEGs they contain, fill color corresponding average correlation between gene expression and plant susceptibility, border color correspond to average LFC upon inoculation. Four major communities are labelled and their number of genes indicated. (E) Distribution of infection LFC and correlation between LFC and susceptibility phenotype for acclimation dependent and independent DEGs. Violin plots show a gaussian kernel, median (dot) and standard deviation (dotted lines).

Next, we performed an analysis of variance on the 9,580 DEGs to determine which of the plant genotype, infection status, acclimation, and their interactions, contributed the most to expression variation for each gene. As expected, infection contributed significantly (Benjamini-Hochberg corrected p-val <1E-3) to the expression variance for 8,523 genes (89%). Genotype and acclimation contributed significantly to the expression variance for 3,111 and 3,716 genes (32.5% and 38.8%) respectively (**Fig 2C, Table S4**). Considering genes the expression variance of which is significantly altered by either acclimation alone or interaction between acclimation and any other factor, acclimation had an impact on the expression of 4,430 genes responsive to infection (46.2% of DEGs, **Fig S3, Table S5**).

To document the relationship between transcriptional response to *S. sclerotiorum* inoculation and acclimation, we built a gene co-expression hierarchical network with genes modulated by inoculation both in the differential and ANOVA analyses. For this, we used normalized read counts to calculate Spearman rank correlation coefficient for all pairwise gene comparisons across our 54 RNA-seq samples. Highly co-expressed gene pairs were grouped into hierarchical gene communities using the HiDeF algorithm (Zheng et al., 2021). 6,620 genes were included into communities of at least four genes (**Fig 2D, Data S1**). Four major top-level communities (labeled α to δ in **Fig 2D**) encompassed 5,933 genes (89.6% of the network). In average, communities α and β included genes with expression anticorrelated with resistance (putative susceptibility factors, **Table S6**), frequently acclimation dependent and downregulated upon *S. sclerotiorum* inoculation and upon heat stress. By contrast, genes from communities and γ and δ had expression correlated with resistance, frequently acclimation-independent, up-regulated upon *S. sclerotiorum* inoculation and heat stress (**Fig 2D, Fig S4**). Accordingly, the median LFC upon inoculation was -1.56 in acclimation-dependent DEGs but 0.70 in acclimation independent DEGs, the median correlation between LFC and susceptibility phenotype was 0.09 in acclimation-dependent DEGs but -0.01 in acclimation independent DEGs (**Fig 2E**). Genes downregulated by infection were 64.3% and 46.7% among genes acclimation-dependent and independent respectively (1.37-fold enrichment). Reciprocally, genes upregulated by infection were 35.5% and 52.6% among genes acclimation-dependent and independent respectively (1.48-fold depletion). Gene the expression of which is correlated (Pearson >0.5) with the susceptibility phenotype were 12.3% and 19.2% among genes acclimation-dependent and independent respectively (1.56-fold enrichment). We conclude that acclimation primarily alters the expression of genes down-regulated by infection and genes associated with disease susceptibility.

### Mediterranean acclimation turns some pathogen-responsive genes into susceptibility factors

To get insights into the role of DEGs in *A. thaliana* QDR against *S. sclerotiorum*, we first analyzed Gene Ontologies (GO) enriched in each of the four major top-level gene communities from our hierarchical network, relative to the rest of the network (**Fig 3A, Table S7**). Community α was enriched in 96 biological process (BP) and 9 molecular function (MF) GO, with ‘Starch metabolism’, ‘Photosynthesis’, ‘Translation’, ‘Primary metabolism’, ‘Chlorophyll binding’, ‘Exopeptidase activity’ and ‘Constituent of ribosome’ among the most enriched, reflecting a general downregulation of energetic functions of the plant cell during infection. Community β was enriched in 18 BP and 4 MF GOs with ‘Regulation of gene expression’, ‘Regulation of metabolic process’, ‘Regulation of developmental process’ and ‘Transcription factor activity’ among the most enriched. Community γ was enriched in 68 BP and 22 MF GOs, with ‘Phytoalexin metabolic process’, ‘Response to chitin’, ‘Chorismate metabolic process’, ‘Immune response’, ‘Glutathione binding’, ‘Oxidoreductase activity’, ‘Carbohydrate binding’ and ‘Ion binding” among the most enriched, reflecting the probable involvement of genes from this community in disease resistance. Finally, community δ was enriched in 40 BP and 10 MF GOs, with ‘Vesicle-mediated transport’, ‘Protein catabolic process’, ‘Response to osmotic stress’ and ‘Signal transduction’ among the most enriched, consistent with a role in stress response.

**Fig 3.**
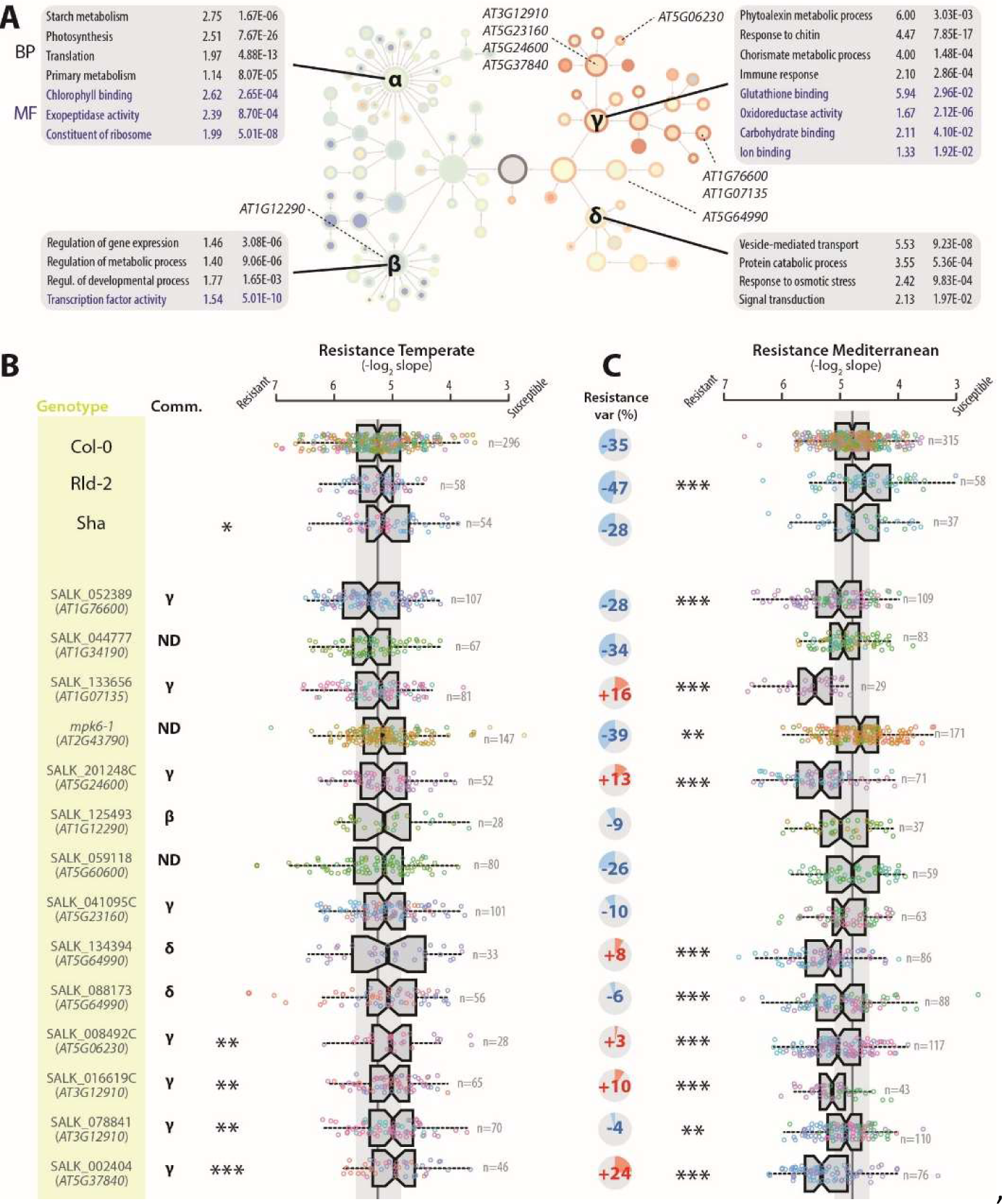
Functional analysis of pathogen-induced genes. (A) Gene ontology (GO) enrichment in the four major gene communities identified based on a co-expression network of genes mis-regulated by *S. sclerotiorum* infection. Communities α, β, γ, δ are labelled on the hierarchical network shown with the same layout as in Fig2D. A selection of the most enriched biological process (BP, black) and molecular function (MF, blue) GOs are labelled, with enrichment fold and adjusted p-value relative to *A. thaliana* genome indicated. Genes analyzed through mutant phenotyping are labeled according to their position in the network. (B, C) Disease resistance phenotype of natural accessions and mutant plants following temperate acclimation (B) or Mediterranean acclimation (C) and inoculated by *S. sclerotiorum*. Comm. Major community of the co-expression network to which the gene belongs. ND, gene not differentially expressed upon *S. sclerotiorum* inoculation (not part of the co-expression network). Pie chart in (C) indicate mean % variation of disease resistance relative to infection following temperate acclimation. Boxplots show first and third quartiles (box), median (thick line), and the most dispersed values within 1.5 times the interquartile range (whiskers). Colors of the data points indicate independent inoculation experiments. Leaves from n=29 to 315 plants were tested for each genotype. Significance of the difference from Col-0 wild type was assessed by a Student’s t test followed by Benjamini-Hochberg correction for multiple testing (*** p<0.01, ** p<0.05., * p<0.1).

To study the role of DEGs in disease resistance against *S. sclerotiorum*, we analyzed the phenotype of 14 mutant lines in the Col-0 background corresponding to 11 distinct genes, with a focus on genes from community γ that were not previously associated with plant immunity (**Table S8**). For comparison purposes, we included mutants in one gene from community β (*AT1G12290*), one from community δ (*AT5G64990*) and three genes not differentially expressed in our RNA-seq experiment (*AT1G34190*, *AT2G43790* and *AT5G60600*). The natural accessions Col-0, Rld-2 and Sha were used as references. After temperate acclimation (**Fig 3B**), four mutant lines were significantly more susceptible than the Col-0 wild type, affecting genes *AT5G06230*, *AT3G12910* and *AT5G37840* from community γ. After Mediterranean acclimation (**Fig 3C, Table S9**), all three natural accessions were more susceptible than after temperate acclimation, consistent with our previous set of experiments (**Fig 1B**). Rld-2 was more strongly affected by Mediterranean acclimation and became significantly more susceptible than Col-0 in these conditions. To our surprise, *mpk6-1* was the only mutant significantly more susceptible than Col-0 after Mediterranean acclimation. Nine mutants were more resistant than wild type after Mediterranean acclimation, covering genes *AT1G76600*, *AT1G07135*, *AT5G24600*, *AT5G06230*, *AT3G12910* and *AT5G37840* from community γ, *AT5G64990* and *AT5G06230* from community δ. While natural accessions had their resistance phenotype reduced by ∼37% in average after Mediterranean compared to temperate acclimation, only four mutant lines showed >25% resistance reduction, including three in genes not differentially expressed upon inoculation (*AT1G34190*, *AT2G43790* and *AT5G60600*) and one from community G (*AT1G76600*). By contrast, six mutants showed increased resistance after Mediterranean compared to temperate acclimation.

Together these results confirm that community γ includes several genes contributing to resistance against *S. sclerotiorum* after temperate acclimation. Mutations in several genes from community γ render plants more resistant than wild type after Mediterranean acclimation, indicating that they act as susceptibility factors in these conditions. Remarkably, AT5G06230, AT3G12910 and AT5G37840 would classify as resistance factors in temperate-acclimated plants but as susceptibility factors in Mediterranean-acclimated plants.

### Acclimation shifts the repertoire of NAC42-L target genes upon *S. sclerotiorum* inoculation

*AT3G12910* encodes a member of the NAC family of transcription factors that includes several regulators of pathogen and abiotic stress response (Nuruzzaman et al., 2013). Its closest homolog in *A. thaliana* genome is *NAC42/JUNGBRUNNEN1* (*AT2G43000*) (Ooka et al., 2003), we will thus refer to *AT3G12910* as *NAC42-Like* (*NAC42-L*) hereafter. *NAC42-L* is strongly induced upon *S. sclerotiorum* inoculation both in plants temperate- (LFC 7.8 p-adj. 3E-08 in Col-0) and Mediterranean-acclimated (LFC 7.5 p-adj. 8E-22 in Col-0). Yet two mutant alleles of *NAC42-L* resulted in lower disease resistance in temperate acclimated plants but enhanced disease resistance in Mediterranean acclimated plants (**Fig 3B**). To study how acclimation alters the activity of NAC42-L at the molecular level, we analyzed the expression of its target genes upon infection in temperate- and Mediterranean-acclimated plants. For this, we first identified targets presumably regulated by NAC42-L by searching for NAC42-L DNA binding motif determined by (O’Malley et al., 2016) in the promoter of *A. thaliana* genes. This identified 2276 potential NAC-L binding sites in 1795 different gene promoters, with a maximum of 5 binding sites per promoter (**Table S10**). Among NAC42-L targets, 394 genes were DEGs upon *S. sclerotiorum* inoculation in temperate- or Mediterranean-acclimated Col-0 plants (**Fig 4B**). There were 227 NAC-L targets (57.6% of NAC42-L target DEGs) uniquely differential following growth under one of the two climates, indicating a significant switch in the regulation of NAC42-L target genes upon infection according to acclimation. The effect of acclimation on the regulation of NAC42-L target genes upon *S. sclerotiorum* inoculation was clearly detectable in the three accessions we analyzed (**Fig 4C**).

**Fig 4.**
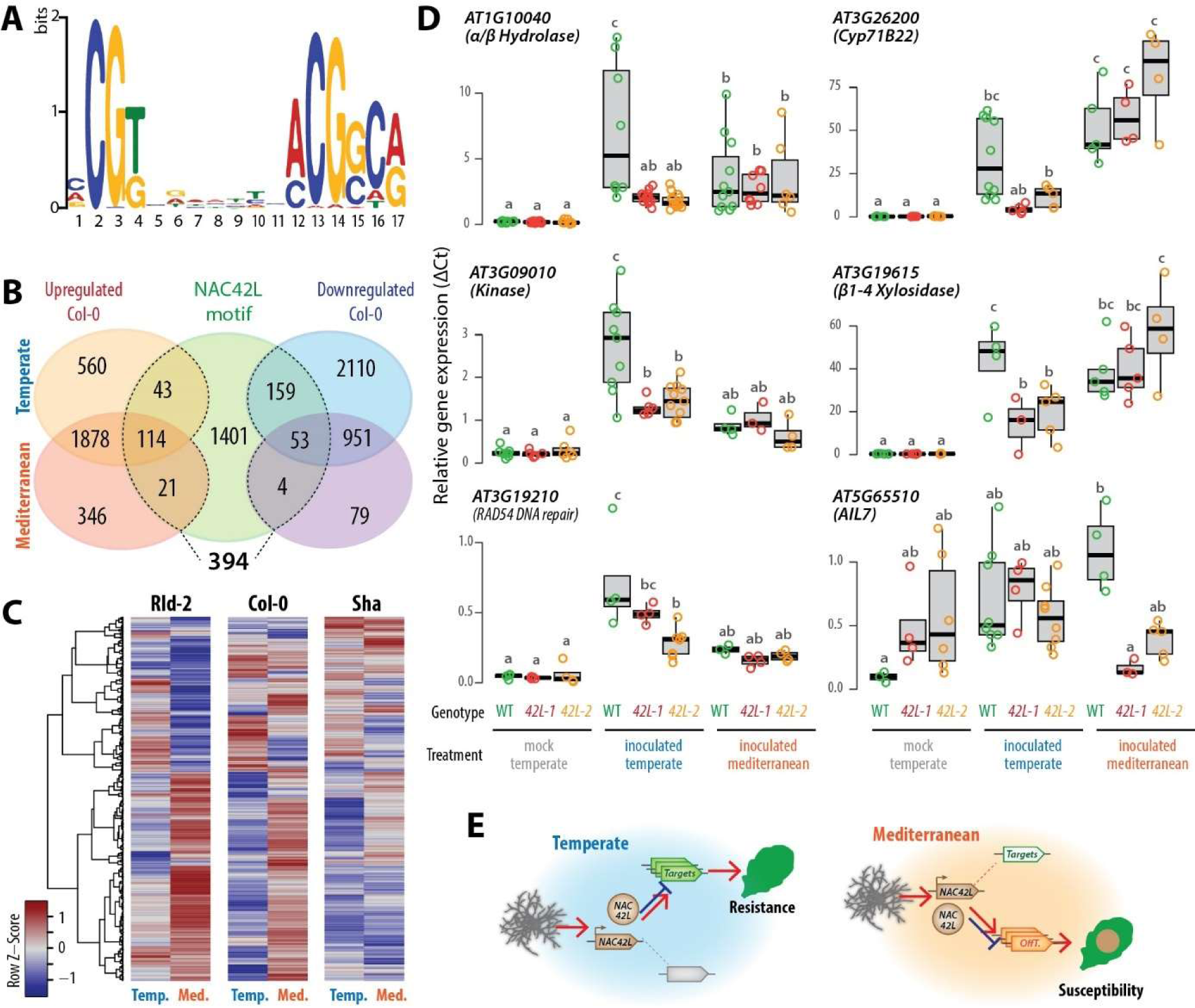
Effect of temperate and Mediterranean acclimation on the regulation of gene expression by the transcription factor NAC42-L. (A) Sequence logo of the promoter motif bound by NAC42-L according to DAP-seq data. (B) Distribution of genes harboring NAC42-L motifs in their promoter between up- and down-regulated genes upon *S. sclerotiorum* inoculation in Col-0 plants temperate- and Mediterranean-acclimated. (C) Relative induction of the 394 genes differentially expressed upon inoculation harboring NAC42-L motifs in their promoter in Col-0 and Sha accessions following temperate (Temp.) and Mediterranean (Med.) acclimation. (D) Expression of six NAC42-L predicted targets in wild type (WT) and *nac42-L* mutants (*42-L1, 42-L2*) in mock-treated and *S. sclerotiorum*-inoculated plants following temperate acclimation and *S. sclerotiorum*-inoculated plants following Mediterranean acclimation. Boxplots show expression independent measurements for 3-9 plants (dots) with first and third quartiles (box), median (thick line), and the most dispersed values within 1.5 times the interquartile range (whiskers). Letters indicate groups of significance determined by a Tuckey HSD test following one-way ANOVA. (E) Schematic representation of the proposed mechanism through which acclimation switches NAC42-L from a positive regulator of disease resistance (temperate acclimation) to a negative regulator (Mediterranean acclimation). *S. sclerotiorum* inoculation triggers the expression of *NAC42-L* (brown arrow) and accumulation of NAC42-L protein (brown circle) which regulates positively (red arrow) or negatively (blue blocked arrow) target genes. Upon temperate acclimation, NAC42-L targets (green arrow) may positively contribute to disease resistance, while upon Mediterranean acclimation, NAC42-L “off-targets” (orange arrows) mostly promote susceptibility to pathogens.

To test whether the acclimation-mediated switch in NAC42-L targets was dependent on NAC42-L, we measured by quantitative RT-PCR the expression of eight of these targets in two *nac42-L* mutant lines (*42-L1* and *42-L2*) following temperate and Mediterranean acclimation (**Fig 4D**, **Table S11, Fig S5**). Six of these genes had an expression significantly altered by the inactivation of *NAC42-L*, supporting their position as targets of NAC42-L regulation. Yet for all of them, the impact of NAC42-L inactivation on their expression was only detected after one particular acclimation regime. Indeed, *AT1G10040*, *AT3G09010*, *AT3G26200* and *AT3G19615* were upregulated upon *S. sclerotiorum* inoculation following temperate acclimation in wild-type plants but significantly less in *42-L1* and *42-L2* plants, while the expression of these genes was similar in all three genotypes following Mediterranean acclimation. Conversely, AT5G65510 showed a similar expression in wild type and *nac42-L* mutant lines upon inoculation following temperate acclimation, but it was significantly mis-regulated in *nac42-L* mutants following Mediterranean acclimation. Together, these results suggest that acclimation alters the contribution of NAC42-L to quantitative disease resistance by switching the repertoire of genes regulated by this transcription factor (**Fig 4E**).

## DISCUSSION

Phenotypic plasticity, a component of acclimation, allows plant species to adjust to environmental conditions, together with adaptation through natural selection or migration to follow conditions to which they are adapted. Understanding the molecular mechanisms of acclimation is crucial for predicting changes in species distributions, community composition and crop productivity under climate change. In this work we show that *A. thaliana* Mediterranean acclimation is detrimental for disease resistance to the fungus *S. sclerotiorum* and converts several genes that contribute positively to quantitative immunity following temperate acclimation into susceptibility factors. Mediterranean acclimation involves a shift in the repertoire of targets of the pathogen-induced transcription factor NAC42-like that may impair the regulation of quantitative immune responses.

Experiments in controlled conditions have been instrumental in unraveling complex stressor interactions through tightly controlled factorial experiments. These studies emphasized that combined effects of various environmental stressors resulted in unique transcriptional changes distinct from individual stress responses (Sewelam et al., 2014; Zandalinas and Mittler, 2022). These interactions can be synergistic, where stressors amplify each other’s negative effects, or antagonistic, where they dampen each other’s impacts (Zarattini et al., 2021). Research on plant-pathogen interactions under abiotic constraints often relies on long-lasting stable temperature shifts, overlooking the complex acclimation processes plants undergo in response to gradual climatic shifts (Aoun et al., 2017; Desaint et al., 2021). Several studies investigated the effect of temperature acclimation by applying a stable temperature shift over a few days prior to a second stress application. For instance, growth of *A. thaliana* for 7 days at 4°C enhanced survival to freezing in a NPR1-dependent manner (Olate et al., 2018), and two-days growth at 30°C rendered plant more susceptible to the bacterial pathogen *Pseudomonas syringae* pv. *tomato* DC3000 when inoculation is performed either at 23°C or 30°C (Huot et al., 2017). Nevertheless, the impact of day-night temperature cycles on subsequent stress response is rarely considered. We have chosen to approximate realistic climate change scenarios by simulating 30-year day and night average temperatures and photoperiods representing three climates of the Köppen-Geiger classification (Peel et al., 2007). Since the 1980s, Mediterranean climates with dry summer (Cs) have gradually replaced areas with temperate climate (Cf) (Cui et al., 2021). Predictions suggest that the Mediterranean (Csa) climate may replace a portion of the continental (Df, Dw, Ds) climates by the end of the century (Beck et al., 2018; Cui et al., 2021). Significant poleward shifts were observed for temperate (C), continental (D), and polar (E) climates with averages of 35.4, 16.2, and 12.6 km.decennia^-1^ (0.32, 0.15, and 0.11° latitude.decennia^-1^ respectively), and are expected to accelerate in the coming decades (Chan and Wu, 2015; Cui et al., 2021). The three selected climates therefore cover a significant part of *A. thaliana* distribution area and reflect the poleward shift of climate zones and associated changes in temperature and day length. Our work revealed a significant loss of quantitative disease resistance upon Mediterranean acclimation in multiple *A. thaliana* accessions, although the daily average temperature was only 0.7°C higher under Mediterranean acclimation (average 15.67°C) than under temperate acclimation (average 14.96°C). Given the current data, we cannot determine whether the observed phenotypic differences are attributable to daytime temperatures, nighttime temperatures, the photoperiod, or a combination of these factors. Nevertheless, our findings indicate that the typical April conditions of the Mediterranean climate zone, expected to expand by the end of the century due to global warming, are detrimental to resistance against *Sclerotinia* diseases. Combined with episodes of high humidity conducive to infection, global warming may therefore increase the incidence of these plant diseases.

Our global transcriptome analyses indicated that the three *A. thaliana* accessions tended to show a higher number of DEGs when acclimated in conditions close to the climate at their area of origin. Col-0, originating from temperate (Cfb, **Fig. S1**) climate area (Somssich, 2019), and Rld-2, originating from continental (Dfb) climate area (Alonso-Blanco et al., 2016) has more upregulated and downregulated genes when acclimated under temperate and continental conditions respectively. Sha, originating from Mediterranean (Csa) climate (Alonso-Blanco et al., 2016) showed more down-regulated following temperate acclimation but more upregulated genes following Mediterranean acclimation. Although this did not reflect at the phenotype level, this transcriptome pattern suggests that *A. thaliana* accessions adapted to their climate of origin to acclimate more efficiently, producing a stronger immune response at the molecular level. It also suggests that mapping to the Col-0 reference genome did not introduce major bias in gene expression quantification in other accessions. Adaptation to environmental change involves variations in allele frequencies within a population’s gene pool over several generations, while acclimation occurs reversibly within an organism’s lifestyle. Genetic variation is crucial for both plastic and adaptive potential (Fox et al., 2019). Reduced genetic variation from positive selection or limited migration can lower phenotypic plasticity. Conversely, plastic traits may become fixed or constitutively expressed through genetic assimilation (Wood et al., 2023).

High genetic variation in natural populations enhances their ability to withstand and adapt to new biotic and abiotic environmental changes, including climate change (Van Kleunen and Fischer, 2005; Nicotra et al., 2010). This genetic variation partly determines the capacity of plants to sense environmental changes and generate plastic responses. For instance, cis-regulatory and epigenetic variation at the *FLOWERING LOCUS C* floral repressor regulating vernalization can aid plant populations in adapting to temperature fluctuations (Hepworth et al., 2020). Yet, the role of selection and whether gene expression plasticity facilitates or hinders adaptation remains a matter of debate (Levis and Pfennig, 2016). Comparative analysis of gene expression in forest and urban populations of *Anolis* lizards showed that rapid parallel regulatory adaptation to urban heat islands primarily resulted from selection for reduced and/or reversed heat-induced plasticity, which is maladaptive in urban thermal conditions (Campbell-Staton et al., 2021). A meta-analysis of reciprocal transplant experiments indicated that adaptation to new environments only leads to genes losing their expression plasticity by genetic assimilation in rare cases (Chen and Zhang, 2024). In agreement, our results suggest that adaptation to their climate of origin maintained high expression plasticity of immunity genes in *A. thaliana* accessions. These insights will be valuable for assessing the adaptive potential of populations in the face of ongoing global climate change.

We identified three genes the inactivation of which in the Col-0 background lead to increased pathogen susceptibility following temperate acclimation but increased resistance following Mediterranean acclimation. Mutation in six other genes resulted in increased resistance following Mediterranean acclimation but no significant phenotype change following temperate acclimation. Conditionally beneficial or neutral mutations, that are deleterious in some environments but beneficial or neutral in others, have been reported in a wide range of organisms including plants (Elena and de Visser, 2003; Anderson et al., 2013). Recombinant inbred lines of the Brassicaceae plant *Boechera stricta* of diverse origin revealed that selection favored local alleles in contrasted environments, and 8.1% of the assessed markers showed evidence for conditional neutrality for the probability of flowering (Anderson et al., 2013). In the perennial grass *Panicum hallii*, an allele of the *FLOWERING LOCUS T-like 9* locus from coastal ecotypes conferred a fitness advantage only in its local habitat but not at the inland site (Weng et al., 2022).

Loss of function alleles contribute to species adaptation (Olson, 1999; Xu and Guo, 2020) and have played an important role in crop domestication (Monroe et al., 2020). Naturally occurring loss of function variants are relatively rare, with an average 57 per genome in *A. thaliana* (Xu et al., 2019) and 18 per genome in soybean (Torkamaneh et al., 2019) but they are found in 19% of soybean genes and 66% of *A. thaliana* genes. Conditionally neutral mutations are sufficient to drive patterns of local adaptation in simulations (Mee and Yeaman, 2019) and can emerge as a compensation to deleterious mutations (Steinberg and Ostermeier, 2024; Farkas et al., 2022). Simulations of long-term evolution in changing environments produced complex gene regulatory networks with an increased rate of beneficial mutations, while a majority of mutations remain neutral (Crombach and Hogeweg, 2008). Patterns of local adaptation in *A. thaliana* (Fournier-Level et al., 2011a), the complexity of quantitative immunity networks (Delplace et al., 2020) and our focus on inoculation up-regulated genes may explain the high proportion of conditionally beneficial loss-of-function we have identified. This finding suggests that targeted gene knockouts may be a promising strategy to improve climate resilience of plant immunity.

We identified *NAC42-L* (*AT3G12910*) as a resistance factor following temperate acclimation but a susceptibility factor following Mediterranean acclimation. Its closest homolog, ANAC042/ JUNGBRUNNEN1 (AT2G43000), was identified as a regulator of camalexin biosynthesis and positive regulator of resistance against the fungus *Alternaria brassicicola* (Saga et al., 2012), longevity (Wu et al., 2012) and tolerance to heat and drought (Ebrahimian-Motlagh et al., 2017; Shahnejat-Bushehri et al., 2012). In addition, exposure to 90min at 37°C enhanced survival of *JUB1* overexpressors to a subsequent treatment at 45°C, compared with WT and *jub1–1* knock-down seedlings (Shahnejat-Bushehri et al., 2012). Molecular changes induced in plants by heat and other environmental signals persist longer than the signals themselves and modifies subsequent responses, phenomenon referred to as somatic environment memory (SEM). In this work, pathogen inoculations were done in standard conditions, indicating that some form of SEM of previous growth conditions had influenced plant immunity. The molecular mechanisms by which SEM mediates the priming of plant-microbe interactions remain largely unknown. Our results implicated a switch in the transcriptional targets of NAC42-L in this process. The underlying molecular bases may include variation in trans, through changes to the composition, stoichiometry and post-transcriptional regulation of protein complexes including NAC42-L, or variation in cis affecting the conformation and accessibility of target gene promoter regions. Recent studies have identified chromatin state modifications as crucial components in the memory of repeated stress events in plants, particularly in response to heat, cold, and drought priming (Balazadeh, 2022; Crisp et al., 2016; Liu et al., 2022). Future investigations will aim at deciphering which molecular mechanisms mediate NAC-L target switch upon acclimation and what controls the duration and breadth of this switch.

Our results highlight rewiring of quantitative immunity gene networks as a key process in acclimation, with adverse consequence to disease resistance under warm Mediterranean-like climates. We show that acclimation can reverse the contribution of genes upregulated by pathogen inoculation to the disease resistance phenotype. We identified several mutations mitigating the negative impact of Mediterranean acclimation on disease resistance, opening perspectives for the preservation of plant immunity functions in a warming climate context.

## MATERIALS AND METHODS

### Plant material and growth conditions

*A. thaliana* Natural accessions and mutant lines were obtained from the Nottingham Arabidopsis Stock Center. We selected Col-0 (CS76778, 6909), Rld-2 (CS78349, 7457) and Shahdara (Sha, CS78397, 6962) as three natural accessions of *A. thaliana* originating from areas with contrasted climate conditions. Plants were grown in jiffy pots for 35 or 70 days in Percival E41-L3 and E41-L2PLT growth cabinets equipped with ultra-sonic humidifier, far-red LED clusters, closed-loop light dimming, Intellus and WeatherEZE controllers. We set day and night temperatures for each climate according to ERA5T models based on 30-year average of hourly weather simulations for daily maximum and minimum temperatures for the month of April at GPS coordinates 56.25°N, 34.19°E (Continental climate, origin of Rld-2 accession); 38.35°N, 68.48°E (Mediterranean climate, origin of Sha accession) and 38.30°N, 92.30°O (Temperate climate) according to https://www.meteoblue.com consulted on April 2017 (**Fig S1**). Day temperatures were 11°C, 20°C and 23°C and night temperatures were 1°C, 9°C and 7°C for continental, temperate and Mediterranean climates respectively. Plants were grown in long day under 190 µmol/m²/s light, with photoperiod variation between climates to represent photoperiod variability during April in the northern hemisphere, water was kept not limiting for the whole experiment at 80% relative humidity. These growth conditions were classified into Continental, Mediterranean and Temperate according to Köppen-Geiger classification (Cui et al., 2021) of climate at the corresponding GPS coordinates. Inoculations were performed on detached leaves at a constant 23°C under constant 40 µmol/m²/s light and high humidity following the procedure described in (Barbacci et al., 2020).

### Fungal strains and disease resistance phenotyping

0.5-cm-wide plugs of PDA agar medium containing *S. sclerotiorum* strain 1980, grown for 72 hours at 20°C on 14 cm Petri dishes, were placed on the adaxial surface of detached leaves. These leaves were positioned in a Navautron system (Barbacci et al., 2020), and records were made using high-definition (HD) cameras “3MP M12 HD 2.8-12mm 1/2.5 IR 1:1.4 CCTV Lens” every 10 minutes. For each genotype, a minimum of 28 leaves were imaged from a minimum of two independent acclimation and inoculation experiments. Kinetics of S. sclerotiorum disease lesions were analyzed using INFEST script v1.0 (https://github.com/A02l01/INFEST). Statistical analyses of disease phenotypes were conducted using the Tukey test or the Student t test followed by Benjamini-Hochberg correction for multiple testing in R 4.2.1. Disease susceptibility (**Fig. 1**) corresponded to the slope of disease lesion growth over time. Resistance (**Fig. 3**) corresponded to -log_2_ of the slope of disease lesion growth over time.

### RNA collection and sequencing

Total RNA was extracted from a 3mm-wide ring of leaf tissue at the edge of ∼1.5cm wide disease lesions collected at 30 hours post inoculation (hpi) for temperate and mediterranean acclimation and 48 hpi for continental acclimation. The samples were harvested with a scalpel on a cool glass slide and immediately frozen in liquid nitrogen. Samples were ground with metal beads (2.5 mm) in a Retschmill apparatus (24hertz for 2x1min). RNA was extracted using the RNAplus kit (Macherey Nagel) following the manufacturer’s instructions. A Turbo DNAse treatment (Ambion) was applied to remove genomic DNA. The quality and concentrations of RNAs preparations were assessed with an Agilent Bioanalyzer using the Agilent RNA 6000 Nano kit. For the analysis of gene expression in natural accessions, libraries synthesis and sequencing was outsourced to Fasteris SA (Plan-les-Ouates, Switzerland). Libraries were sequenced as paired-end reads on an Illumina HiSeq 2500 instrument in High Output v4 mode with 2x125+8 cycles on 7 lanes of HiSeq Flow Cells v4 with the HiSeq SBS Kit v4. Basecalling was performed with the HiSeq Control Software 2.2.58, RTA 1.18.64.0 and CASAVA-1.8.2. Reads QC was performed using spiked-PhiX in-lane controls yielding Q30 error rate <0.4% for all lanes. Paired-end reads were trimmed and mapped to the TAIR10.0 reference genome using the RNA-seq analysis tool of the CLC Genomics Workbench 11.0.1 software (Qiagen). The following mapping parameters were used: mismatch cost 2, insertion cost 3, deletion cost 3, length fraction 0.8, similarity fraction 0.8, both strands mapping, and 10 hits maximum per read, with expression value given as total read count per gene.

### Differential expression and expression variance analyses

Differential gene expression analysis was performed with the DESeq2 Bioconductor package version 1.8.2 (Love et al., 2014) in R 3.4.0 in a pairwise manner using expression in uninfected plants as a reference with ∼replicates + inoculation as the design formula. Genes with baseMean 0 in all differential comparisons and non-nuclear genes were discarded from further analyses. Genes with |Log2 Fold Change|≥2 and Bonferroni-adjusted p-val<0.001 in DESeq2 Wald test were considered significant for differential expression. For ANOVA, read counts were mean-normalized to homogenize the total number of mapped reads per sample. The ANOVA was performed on each gene using the dplyr package in R with ReadCount ∼ Genotype * Infection * Climate as the model formula. P-values associated with each factor were corrected for multiple testing using the Benjamini-Hochberg procedure.

### Gene network reconstruction and analyses

For gene network reconstruction, we focused on the 7,279 genes whose expression was pathogen inoculation-dependent both in the differential analysis (|Log2 Fold Change|≥2 and Bonferroni-adjusted p-val<0.001) and in the ANOVA analysis (Benjamini-Hochberg corrected p-val <1E-7). Pairwise Spearman rank correlation was calculated for the expression of these genes using the rcorr function from the R package Hmisc, using the raw counts per gene from all 54 RNA-seq samples. The top 25 correlated genes (Spearman ρ>0.85) was extracted for each gene. The top 25 expression correlations were used as edges for network reconstruction with weight ρ. Reconstruction of the hierarchical gene cluster network was performed using the Community Detection 1.12.0 plugin in Cytoscape 3.10.0, using the HiDeF algorithm with maximum resolution 45.0, consensus threshold 65, persistent threshold 6 and the Louvain algorithm. Correlation with susceptibility was the Spearman rank correlation coefficient between normalized read counts (averaged over three replicates) for each gene and the slope of disease lesion growth. Average LFC was the mean log2 fold change over all nine genotype-acclimation modalities tested. Values for gene clusters are the mean of values for all genes in a cluster. Gene ontology enrichment were analyzed with the BinGO plugin in Cytoscape 3.10.0 using a hypergeometric test with Benjamini and Hochberg false discovery rate correction, at significance level 0.05 with *A. thaliana* whole annotation as a reference set.

### Characterization of *A. thaliana* mutant lines

T-DNA insertion lines in AT1G76600 (SALK_052389), AT1G34190 (SALK_044777), AT1G07135 (SALK_133656), AT2G43790 (mpk6-1), AT5G24600 (SALK_201248C), AT1G12290 (SALK_125493), AT5G60600 (SALK_059118), AT5G23160 (SALK_041095C), AT5G64990 (SALK_088173), AT5G06230 (SALK_008492C), AT3G12910 (SALK_016619C and SALK_078841) and AT5G37840 (SALK_002404) in the Col-0 background were obtained from the Nottingham Arabidopsis Stock Centre. To identify homozygous insertion lines, the lines were genotyped by PCR and, if needed, self-crossed and the progeny genotyped by PCR. The disease resistance phenotype of mutant lines was analyzed as previously described in a total of 23 independent inoculation experiments each including the Col-0 reference, with a minimum of 2 independent experiments for each mutant line. Primers used in all experiments are shown in **Table S12**.

### Identification of NAC42-L predicted target genes

The HMM model NAC_tnt.AT3G12910_col_a_m1 for the unique DAP-seq motif bound by AT3G12910 was obtained from the Plant Cistrome Database (http://neomorph.salk.edu/dap_web/) (O’Malley et al., 2016). *A.thaliana* genes harboring the corresponding motif were identified using FIMO v.5.5.5 (Grant et al., 2011) using the sequence 1Kbp upstream of the translation start site from the Araport11 annotation with 1E-4 as p-value threshold, both strands scanning and NRDB frequencies as a background model

### Quantitative RT-PCR Analyses

RNA for qRT-PCR analysis was extracted from plants 24 hours post-inoculation with *S. sclerotiorum* as for RNA-sequencing. For cDNA synthesis, 1 µg of RNA and 0.5 µL of Transcriptor reverse transcriptase (Roche) were used in a 20 µL reaction volume according to the manufacturer’s protocol. The resulting cDNA diluted 1:10 served as the template for quantitative RT-PCR. qRT-PCR reactions were carried out with 5 pmol of specific oligonucleotides (**Table S12**), 2 µL of cDNA, and 3.5 µL of SYBR GREEN I in a total volume of 7 µL. Amplification reactions were performed using a LightCycler 480 (Roche Diagnostics) with the following protocol: 9 minutes at 95°C, followed by 45 cycles of 5 seconds at 95°C, 10 seconds at 65°C, and 20 seconds at 72°C. Relative gene expression was calculated as the ratio of target gene expression to the reference gene *AT2G28390* and expressed as the difference between target and reference crossing times (ΔCt).

## ACKNOWLEDGEMENTS

This work was supported by the French Laboratory of Excellence project ‘TULIP’ (ANR-10-LABX-41; ANR-11-IDEX-0002-02), a Starting grant from the European Research Council (ERC-StG-336808), l’Agence Nationale pour la Recherche (ANR-19-CE20-15, ANR-21-CE20-10, ANR-21-CE20-30), and the INRAE. M.D. benefited from a phD grant of the INRAE SPE division. We are grateful to the LIPME Bioinformatics team for invaluable assistance with data storage and analysis. We thank Mehdi Khafif for excellent technical help. ChatGPT-4 and Perplexity AI beta v0 were used to polish some sections of the introduction and discussion of this manuscript.

## DATA AVAILABILITY

Raw RNA-seq reads data and processed gene expression files generated in this work are available under NCBI GEO accession number GSE272240.

## AUTHOR CONTRIBUTIONS

A.B. and S.R. designed research; Natural accessions phenotyping: M.D., J.S.; RNA-seq sampling: J.S.; RNA-seq data analysis: J.S., M.D., S.R.; Analysis of mutant lines genotype and phenotype: M.D; NAC42-L targets identification: M.D; Sampling for qRT-PCR: M.D.; Performed qRT-PCR: M.D., P.C-S, M.Z; Analyzed qRT-PCR results: M.D, P.C-S, SR; Supervised and coordinated research: A.B, S.R; Wrote the paper: M.D and S.R.

## SUPPLEMENTARY MATERIAL

**Supplementary Figure S1.**
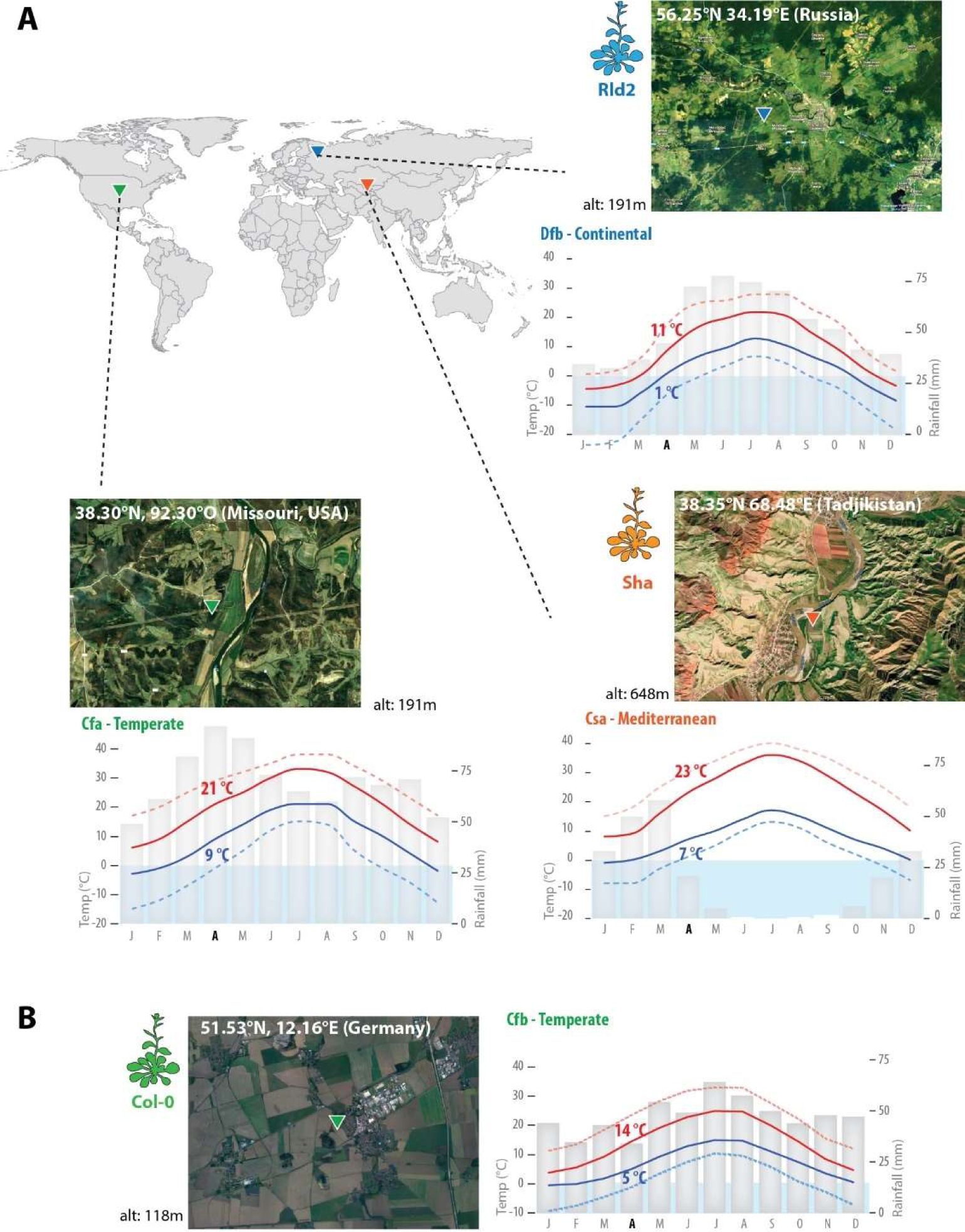
Climate data at sites in the distribution range of *A. thaliana* and *S. sclerotiorum* used for simulated climates in our experiments (A) and at the site of Col-0 accession origin (B). Data come from an ERA5T model of 30-year average of hourly simulations collected from meteoblue.com. Satellite views of the designated coordinates were obtained from Google Maps. Grey bars show monthly precipitations in mm, red lines are mean daily maximum (plain) and hot days maximum (dotted), blue lines are mean daily minimum (plain) and cold nights minimum (dotted). Values for April are labelled. The corresponding Köppen-Geiger climate was obtained from climate-data.org and coded as follows: Csa, hot summer Mediterranean climate; Cfa, humid subtropical climate; Cfb, Temperate oceanic climate or subtropical highland climate; Dfa, Hot-summer humid continental climate. Alt., altitude; Temp., Temperature.

**Supplementary Figure S2.**
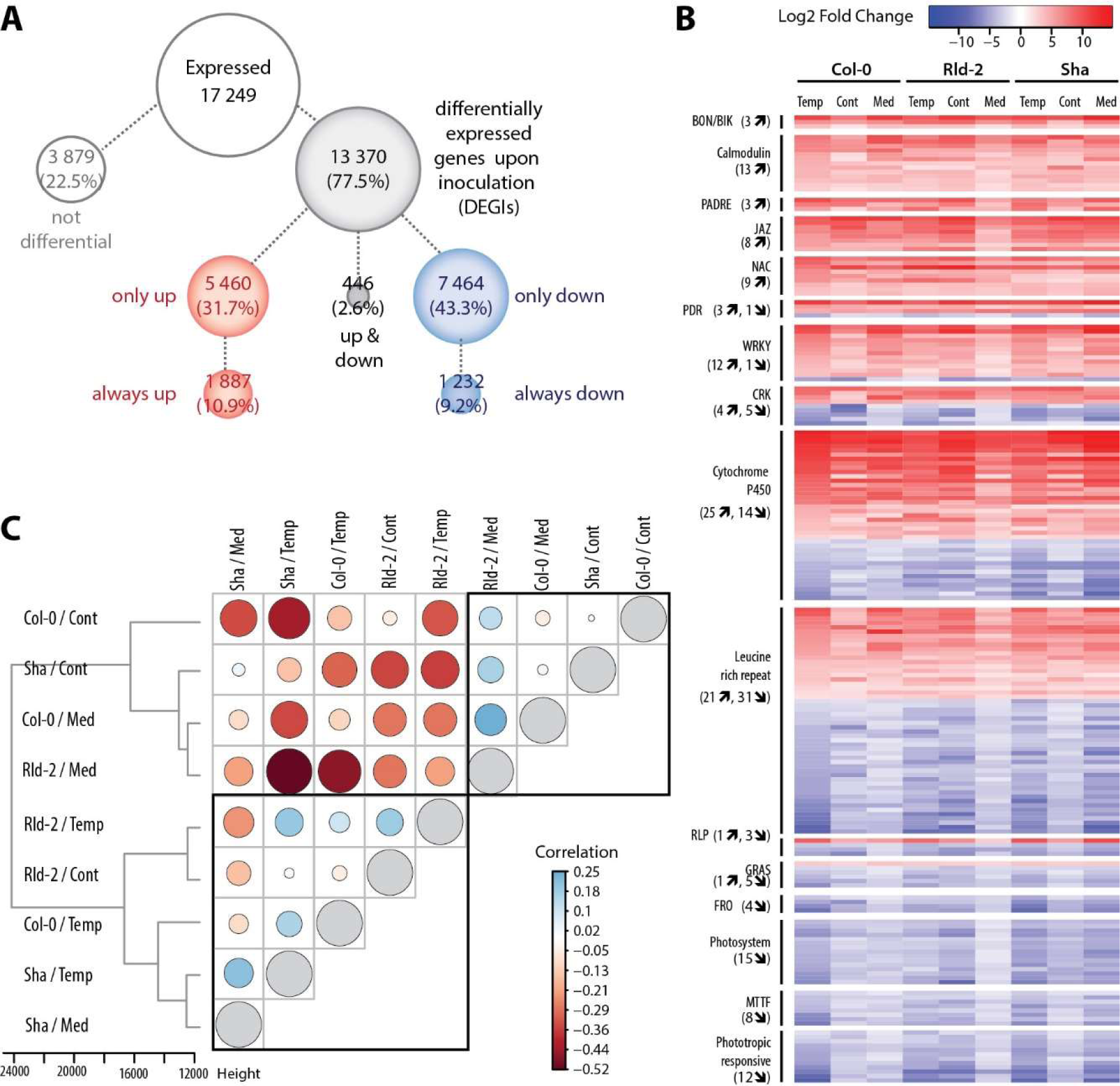
*Arabidopsis* genes differentially expressed upon *S. sclerotiorum* inoculation analyzed with relaxed thresholds. Analysis of <u>g</u>enes differentially expressed upon inoculation at |Log2 Fold Change|≥1.5, adjusted p-val<0.1, using non-inoculated plants as reference in each of nine conditions (three climate priming, times three plant genotypes). **(A)** Identification of 13 370 differentially expressed genes (DEGs) upon inoculation. In conditions where they are differential, 5 460 DEGs were upregulated only, 7 464 were down-regulated only, and 446 were either up or down-regulated. We detected 1 887 DEGs upregulated in all nine conditions and 1 232 DEGs down-regulated in all nine conditions, representing 10.9% and 9.2% of the expressed genes respectively. These 3 119 genes are differentially expressed in a consistent manner regardless of plant genotype and acclimation, they can therefore be regarded as a core transcriptome responsive to *S. sclerotiorum* in *A. thaliana*. **(B)** Heatmap of Log2 fold change for 200 genes forming major functional groups including DEGs always up and always down (numbers indicated between brackets). Major functional groups in the core transcriptome included the PRR-associated *BOTRYTIS-INDUCED KINASE 1* (*BIK1*), *BONZAI1-ASSOCIATED PROTEIN (BAP) 1* and *2*, members of the Calmodulin (CaM) and CaM-binding, the pathogen and abiotic stress response, cadmium tolerance, disordered region-containing (PADRE), the jasmonate-zim-domain proteins (JAZ), and the NAC-domain transcription factor families, with all core DEGs being upregulated by *S. sclerotiorum* inoculation. Ferric reduction oxidases (FRO), Mitochondrial transcription termination factors (MTTF), Photosystems I and II, phototropin and phototropic-responsive NPH3 genes showed all core DEGs downregulated. The pleiotropic drug resistance (PDR), cysteine-rich receptor-like protein kinase (CRK), WRKY and GRAS transcription factor, cytochrome P450, Leucine-rich repeat (LRR), receptor like protein (RLP) families included several core genes responsive to *S. sclerotiorum* either consistently up- or down-regulated. **(C)** Distance tree and correlation matrix showing the similarity in the 13 370 DEGs regulation across conditions. Tree based on Manhattan distance between samples and Ward clustering, correlation values shown by bubbles are Spearman rank correlations calculated using LFC for 13 370 DEGs in each condition. Transcriptomes measured after priming under temperate acclimation clustered together and with the transcriptome of Rld-2 primed under continental acclimation and that of Sha under Mediterranean acclimation. The remaining four conditions formed a second cluster. There was no clear clustering based on genotype or climates alone, suggesting a significant interaction between these factors. Cont, continental; Temp, temperate; Med, Mediterranean.

**Supplementary Figure S3.**
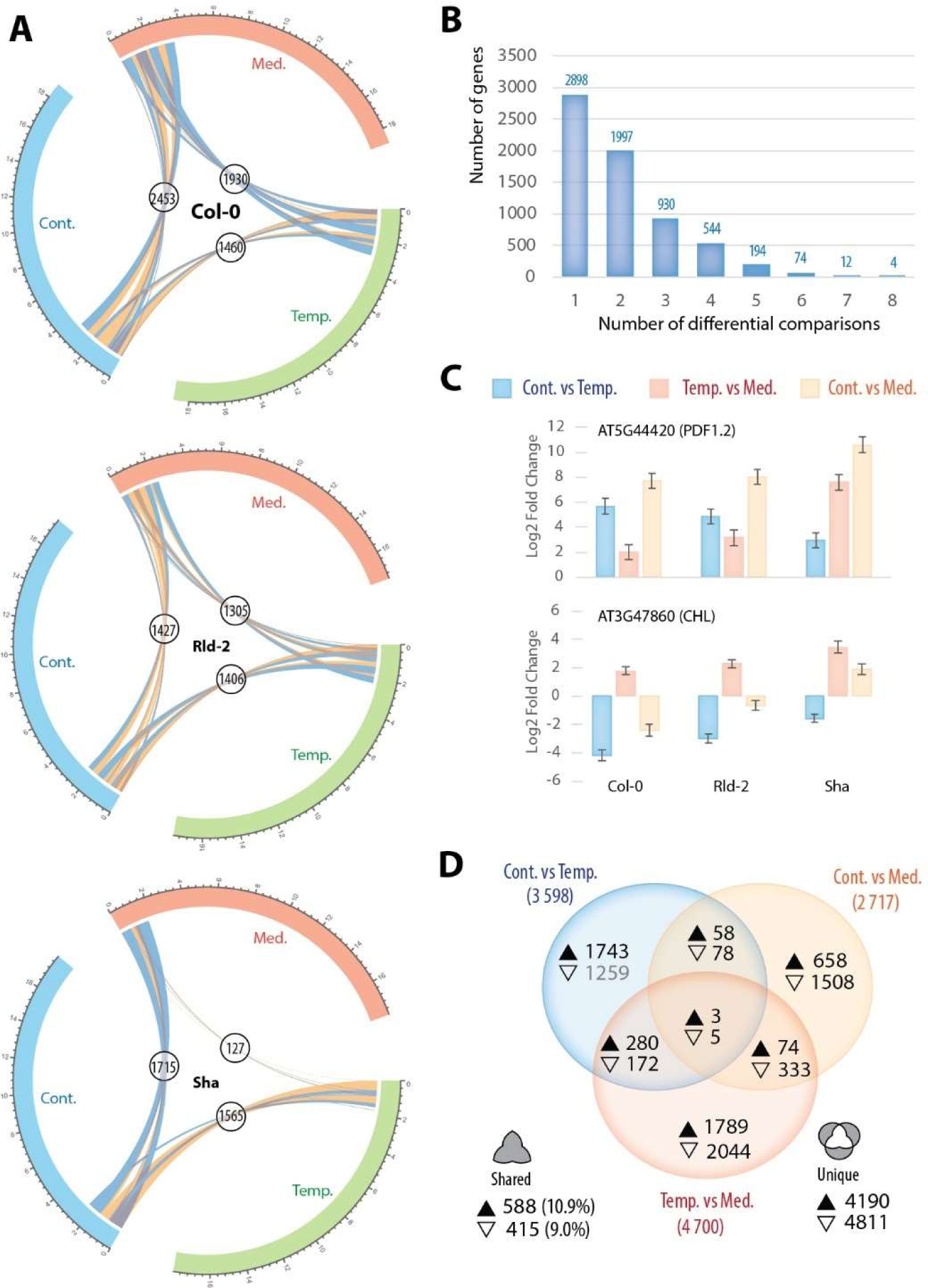
*A. thaliana* genes differentially expressed in pairwise acclimation comparisons. **(A)** Differentially expressed <u>g</u>enes in pairwise comparisons between acclimation regimes for *S. sclerotiorum*-inoculated samples (|LFC|≥1.5, adjusted p-val<0.1). The circos tracks show number of expressed genes (x 1,000) with connectors representing genes differentially expressed between two acclimation regimes (down-regulated in blue, upregulated in yellow). Labels show the sum of up- and down-regulated genes for each acclimation comparison. The number of DEGs ranged from 127 in Sha when comparing temperate and Mediterranean acclimation to 2,453 in Col-0 when comparing continental and Mediterranean acclimation, representing 0.71% to 13% of expressed genes. **(B)** Overall, 6,653 genes were differential in at least one comparison (35.2% of expressed genes). Of those, 4,895 (73.6%) were differentially expressed in one or two acclimation comparisons only, consistent with a specific effect of acclimation on *A. thaliana* transcriptome, and supporting a significant interaction between genotype and acclimation. **(C)** The genes most sensitive to acclimation were four genes differentially regulated in eight out of nine comparisons. These genes encoded plant defensins PDF1.3 (AT2G26010) and PDF1.2 (AT5G44420), the MYB transcription factor LHY (LATE ELONGATED HYPOCOTYL, AT1G01060) and the chloroplastic lipocalin gene CHL (AT3G47860). Histograms show Log2 Fold change of expression for AT5G44420 and AT3G47860 in 3 acclimation comparisons for 3 genotypes. Error bars show estimated standard errors for the estimated coefficients on the log2 scale from DESeq2. **(D)** Venn diagram showing the distribution of DEGs for all three accessions according to the acclimation regimes compared. The full upward triangle indicates up-regulated genes, the empty downward triangle indicates down-regulated genes. For each acclimation comparison the total number of DEGs is indicated between parenthesis. The percentage of shared genes is given relative to the total number of DEGs in acclimation comparisons. Cont., continental acclimation; Med., Mediterranean acclimation; Temp., temperate acclimation.

**Supplementary Figure S4.**
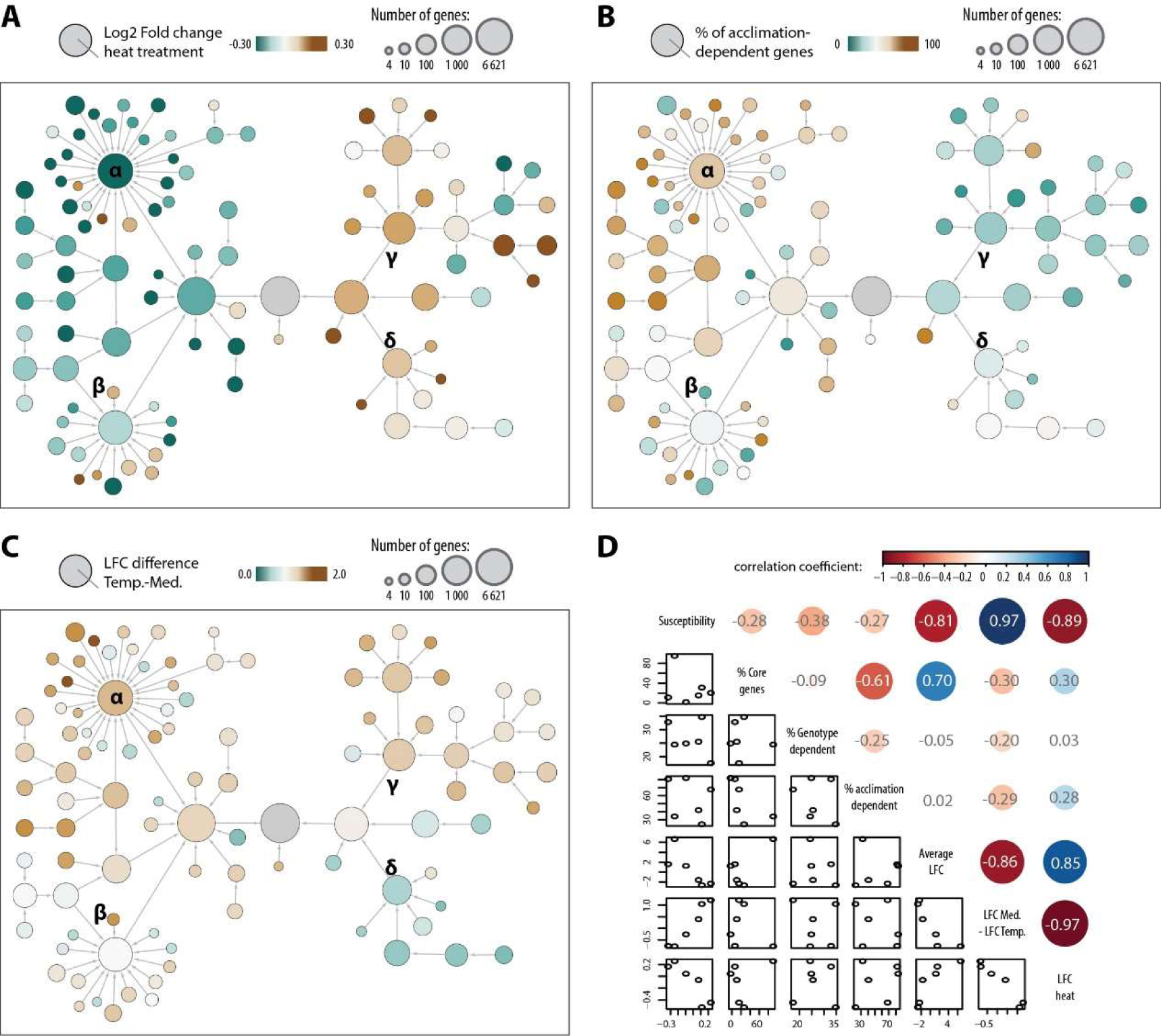
Properties of gene communities in a network of DEGs upon S. sclerotiorum inoculation. **(A)** We mapped the average gene LFC upon heat treatment reported in a recent meta-analysis (Guo *et al*. 2021). Genes from communities γ and δ showed a trend for up-regulation upon heat treatment (average LFC 0.26 and 0.17 respectively) while genes from communities α and β were rather down-regulated (average LFC -0.52 and -0.45 respectively). **(B)** We mapped the percentage of gene from each community considered acclimation dependent based on our ANOVA analysis. Communities γ and δ had a low proportion of acclimation-dependent genes while community α had a majority of acclimation-dependent genes. **(C)** Considering the clear phenotypic effect of Mediterranean acclimation, which had the highest day temperature and highest daily thermal amplitude, we mapped the average LFC variation between plants acclimated under temperate and Mediterranean climates. Average LFC variation was >1.0 for communities α and γ but <-0.7 for community δ. (D) To summarize these analyses, we calculated the correlation between properties of the six largest gene communities. We observed a clear correlation between association with susceptibility phenotype and LFC variation upon temperate and Mediterranean acclimation (0.97), and anti-correlation with average LFC upon *S. sclerotiorum* inoculation (-0.81 and -0.86). This suggested that in our experiments, differential gene expression mostly associated with a decrease in plant susceptibility which is strongly altered upon Mediterranean acclimation. Reference: Guo, M., Liu, X., Wang, J., Jiang, Y., Yu, J., & Gao, J. (2021). Transcriptome profiling revealed heat stress-responsive genes in *Arabidopsis* through integrated bioinformatics analysis. *Journal of Plant Interactions*, *17*(1), 85–95. https://doi.org/10.1080/17429145.2021.2014580

**Supplementary Figure S5.**
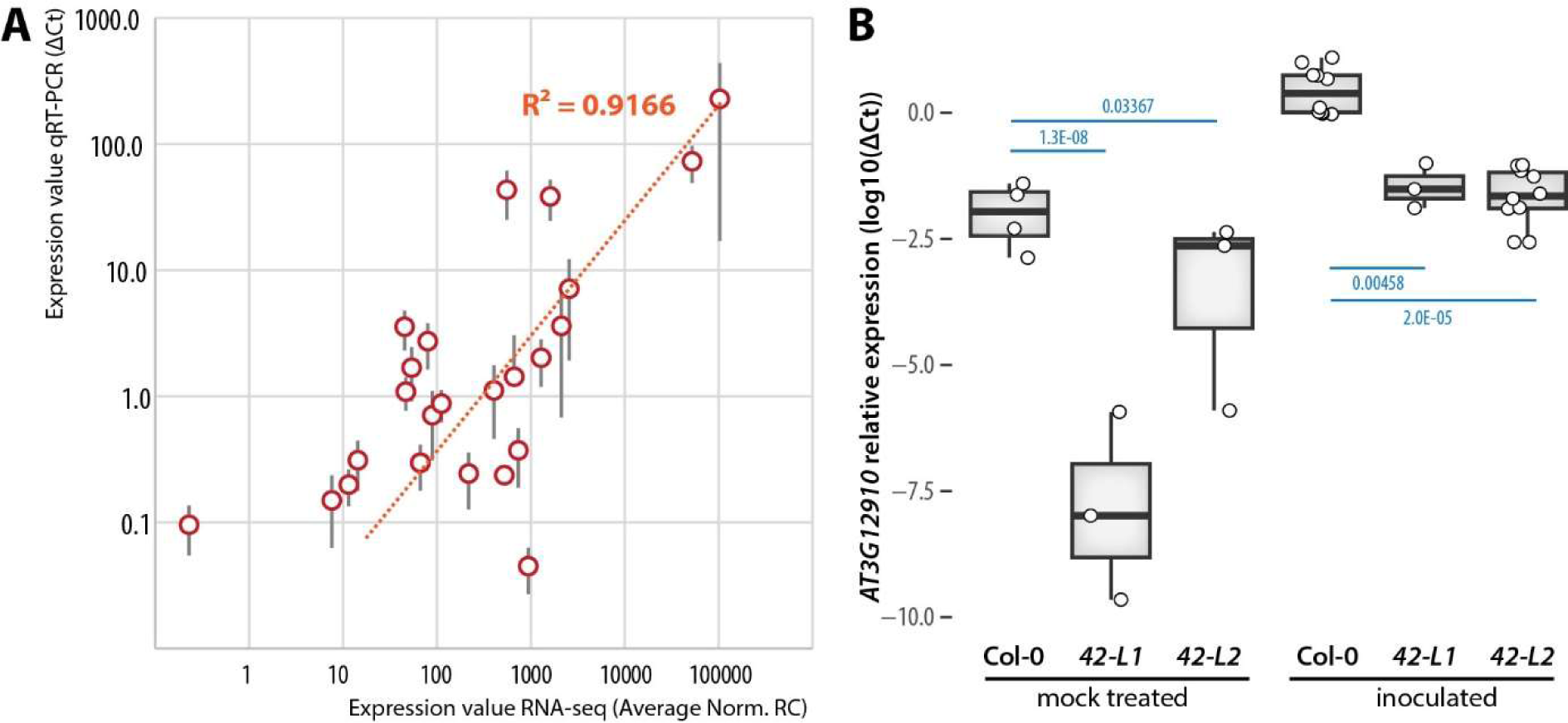
Analysis of gene expression by quantitative RT-PCR in wild type and NAC42-L mutant lines. **(A)** Relationship between gene expression in Col-0 determined by RNA-sequencing (X axis) and quantitative RT-PCR (Y axis). Error bars show standard error of the mean from 3 to 11 independent replicates. Dots represent expression of AT1G10040, AT1G56130, AT3G09010, AT3G12910, AT3G19210, AT3G19615, AT3G26200, AT4G39950, AT5G65510 in mock-treated and S. sclerotiorum inoculated plants grown under temperate and Mediterranean acclimation. (B) Relative expression of NAC42-L (AT3G12910) in Col-0, nac42-L1 and nac42-L2 mutant lines determined by quantitative RT-PCR. Boxplots show expression independent measurements for 3-9 plants (dots) with first and third quartiles (box), median (thick line), and the most dispersed values within 1.5 times the interquartile range (whiskers). P-values were determined by a Student t test with Benjamini-Hochberg correction for multiple testing.

## Supplementary tables online

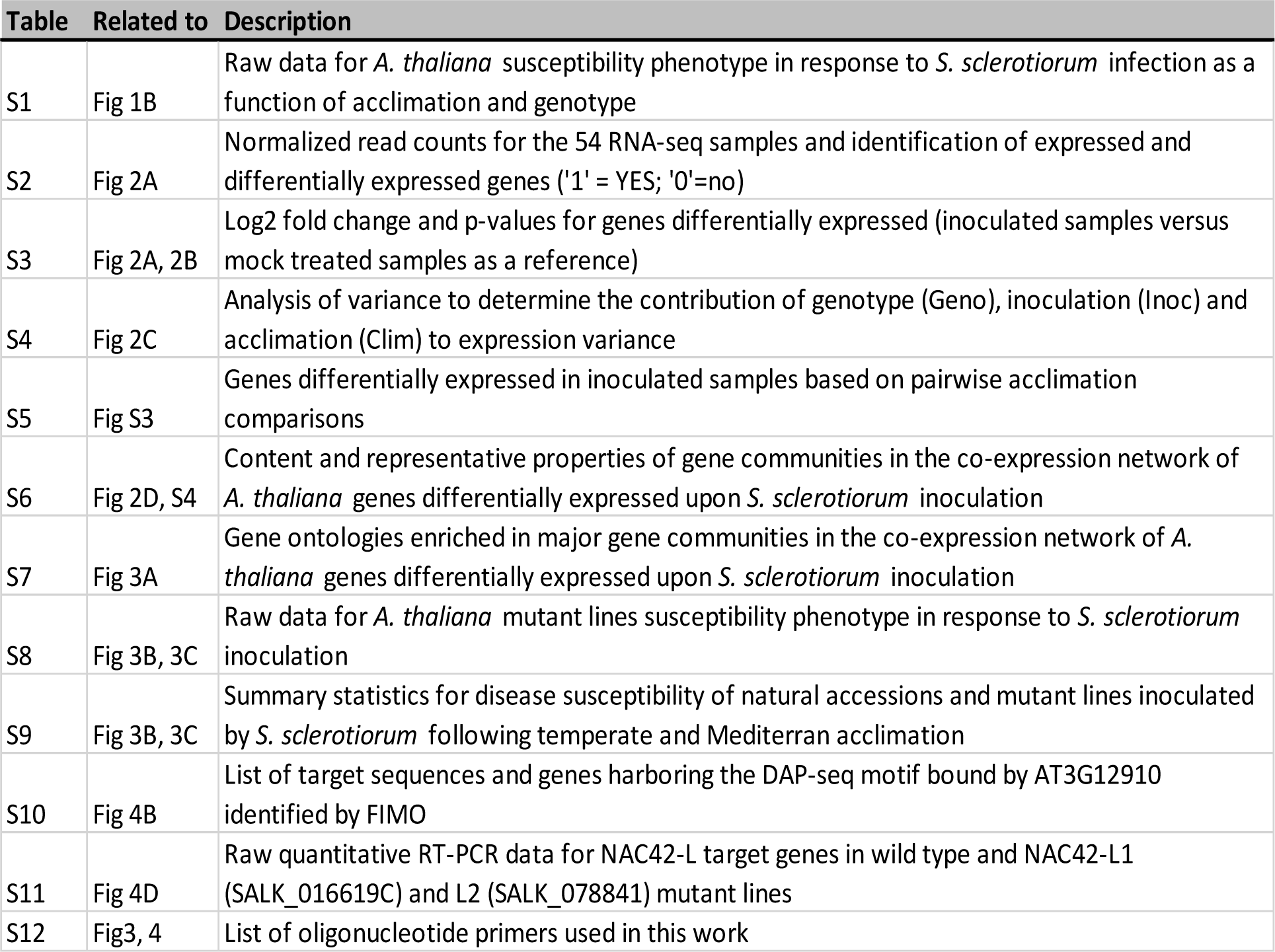

## Supplementary Data online

**Data S1**. Cytoscape session file containing the network of *A. thaliana* genes differentially expressed upon *S. sclerotiorum* inoculation, associated metadata and hierarchical clustering network.

## Notes

https://www.ncbi.nlm.nih.gov/geo/

### Competing Interest Statement

The authors have declared no competing interest.

## REFERENCES

Alonso-Blanco, C. et al. (2016). 1,135 Genomes Reveal the Global Pattern of Polymorphism in Arabidopsis thaliana. Cell 166: 481–491.

Anderson, J.T., Lee, C.-R., Rushworth, C.A., Colautti, R.I., and Mitchell-Olds, T. (2013). Genetic trade-offs and conditional neutrality contribute to local adaptation. Mol. Ecol. 22: 699–708.

Aoun, N., Tauleigne, L., Lonjon, F., Deslandes, L., Vailleau, F., Roux, F., and Berthomé, R. (2017). Quantitative disease resistance under elevated temperature: Genetic basis of new resistance mechanisms to ralstonia solanacearum. Front. Plant Sci. 8: 1–16.

Badet, T., Léger, O., Barascud, M., Voisin, D., Sadon, P., Vincent, R., Le Ru, A., Balagué, C., Roby, D., and Raffaele, S. (2019). Expression polymorphism at the *ARPC 4* locus links the actin cytoskeleton with quantitative disease resistance to *Sclerotinia sclerotiorum* in *Arabidopsis thaliana*. New Phytol. 222: 480–496.

Balazadeh, S. (2022). A ‘hot’ cocktail: The multiple layers of thermomemory in plants. Curr. Opin. Plant Biol. 65: 102147.

Barbacci, A., Navaud, O., Mbengue, M., Barascud, M., Godiard, L., Khafif, M., Lacaze, A., and Raffaele, S. (2020). Rapid identification of an Arabidopsis NLR gene as a candidate conferring susceptibility to Sclerotinia sclerotiorum using time-resolved automated phenotyping. Plant J. 103: 903–917.

Bebber, D.P., Holmes, T., and Gurr, S.J. (2014). The global spread of crop pests and pathogens. Glob. Ecol. Biogeogr. 23: 1398–1407.

Bebber, D.P., Ramotowski, M.A.T., and Gurr, S.J. (2013). Crop pests and pathogens move polewards in a warming world. Nat. Clim. Chang. 3: 985–988.

Beck, H.E., Zimmermann, N.E., McVicar, T.R., Vergopolan, N., Berg, A., and Wood, E.F. (2018). Present and future Köppen-Geiger climate classification maps at 1-km resolution. Sci. Data 5: 180214.

Bernard, F., Chelle, M., Fortineau, A., Riahi El Kamel, O., Pincebourde, S., Sache, I., and Suffert, F. (2022). Daily fluctuations in leaf temperature modulate the development of a foliar pathogen. Agric. For. Meteorol. 322: 109031.

Bohn, L., Huang, J., Weidig, S., Yang, Z., Heidersberger, C., Genty, B., Falter-Braun, P., Christmann, A., and Grill, E. (2024). The temperature sensor TWA1 is required for thermotolerance in Arabidopsis. Nature.

Brancalion, P.H.S., Oliveira, G.C.X., Zucchi, M.I., Novello, M., van Melis, J., Zocchi, S.S., Chazdon, R.L., and Rodrigues, R.R. (2018). Phenotypic plasticity and local adaptation favor range expansion of a Neotropical palm. Ecol. Evol. 8: 7462–7475.

Burghardt, L.T., Runcie, D.E., Wilczek, A.M., Cooper, M.D., Roe, J.L., Welch, S.M., and Schmitt, J. (2016). Fluctuating, warm temperatures decrease the effect of a key floral repressor on flowering time in Arabidopsis thaliana. New Phytol. 210: 564–576.

Campbell-Staton, S.C., Velotta, J.P., and Winchell, K.M. (2021). Selection on adaptive and maladaptive gene expression plasticity during thermal adaptation to urban heat islands. Nat. Commun. 12: 6195.

Chaloner, T.M., Gurr, S.J., and Bebber, D.P. (2021). Plant pathogen infection risk tracks global crop yields under climate change. Nat. Clim. Chang. 11: 710–715.

Chan, D. and Wu, Q. (2015). Significant anthropogenic-induced changes of climate classes since 1950. Sci. Rep. 5: 13487.

Charng, Y.Y., Mitra, S., and Yu, S.J. (2023). Maintenance of abiotic stress memory in plants: Lessons learned from heat acclimation. Plant Cell 35: 187–200.

Chen, P. and Zhang, J. (2024). Transcriptomic analysis reveals the rareness of genetic assimilation of gene expression in environmental adaptations. Sci. Adv. 9: eadi3053.

Clauw, P., Kerdaffrec, E., Gunis, J., Reichardt-Gomez, I., Nizhynska, V., Koemeda, S., Jez, J., and Nordborg, M. (2022). Locally adaptive temperature response of vegetative growth in Arabidopsis thaliana. Elife 11: 1–17.

Cohen, S.D. (2023). Estimating the Climate Niche of Sclerotinia sclerotiorum Using Maximum Entropy Modeling. J. Fungi 9.

Coolen, S. et al. (2016). Transcriptome dynamics of Arabidopsis during sequential biotic and abiotic stresses. Plant J. 86: 249–67.

Corwin, J.A., Copeland, D., Feusier, J., Subedy, A., Eshbaugh, R., Palmer, C., Maloof, J., and Kliebenstein, D.J. (2016). The Quantitative Basis of the Arabidopsis Innate Immune System to Endemic Pathogens Depends on Pathogen Genetics. PLoS Genet. 12: 1–29.

Crisp, P.A., Ganguly, D., Eichten, S.R., Borevitz, J.O., and Pogson, B.J. (2016). Reconsidering plant memory: Intersections between stress recovery, RNA turnover, and epigenetics. Sci. Adv. 2.

Crombach, A. and Hogeweg, P. (2008). Evolution of Evolvability in Gene Regulatory Networks. PLOS Comput. Biol. 4: e1000112.

Cui, D., Liang, S., and Wang, D. (2021). Observed and projected changes in global climate zones based on Köppen climate classification. WIREs Clim. Chang. 12: e701.

Delplace, F., Huard-Chauveau, C., Dubiella, U., Khafif, M., Alvarez, E., Langin, G., Roux, F., Peyraud, R., and Roby, D. (2020). Robustness of plant quantitative disease resistance is provided by a decentralized immune network. Proc. Natl. Acad. Sci. 117: 18099–18109.

Deng, Y., Bossdorf, O., and Scheepens, J.F. (2021). Transgenerational effects of temperature fluctuations in Arabidopsis thaliana. AoB Plants 13: 1–11.

Derbyshire, M.C. and Raffaele, S. (2023). Till death do us pair: Co-evolution of plant–necrotroph interactions. Curr. Opin. Plant Biol. 76: 102457.

Desaint, H., Aoun, N., Deslandes, L., Vailleau, F., Roux, F., and Berthomé, R. (2021). Fight hard or die trying: when plants face pathogens under heat stress. New Phytol. 229: 712–734.

Ebrahimian-Motlagh, S., Ribone, P.A., Thirumalaikumar, V.P., Allu, A.D., Chan, R.L., Mueller-Roeber, B., and Balazadeh, S. (2017). JUNGBRUNNEN1 Confers Drought Tolerance Downstream of the HD-Zip I Transcription Factor AtHB13. Front. Plant Sci. 8: 2118.

Elena, S.F. and de Visser, J.A.G.M. (2003). Environmental stress and the effects of mutation. J. Biol. 2: 12.

Farkas, Z., Kovács, K., Sarkadi, Z., Kalapis, D., Fekete, G., Birtyik, F., Ayaydin, F., Molnár, C., Horváth, P., Pál, C., and Papp, B. (2022). Gene loss and compensatory evolution promotes the emergence of morphological novelties in budding yeast. Nat. Ecol. Evol. 6: 763–773.

Fournier-Level, A., Korte, A., Cooper, M.D., Nordborg, M., Schmitt, J., and Wilczek, A.M. (2011a). A map of local adaptation in Arabidopsis thaliana. Science 334: 86–9.

Fournier-Level, A., Korte, A., Cooper, M.D., Nordborg, M., Schmitt, J., and Wilczek, A.M. (2011b). A map of local adaptation in Arabidopsis thaliana. Science (80-.). 334: 86–89.

Fox, R.J., Donelson, J.M., Schunter, C., Ravasi, T., and Gaitán-Espitia, J.D. (2019). Beyond buying time: the role of plasticity in phenotypic adaptation to rapid environmental change. Philos. Trans. R. Soc. B Biol. Sci. 374: 20180174.

Garcia-Molina, A., Kleine, T., Schneider, K., Mühlhaus, T., Lehmann, M., and Leister, D. (2020). Translational Components Contribute to Acclimation Responses to High Light, Heat, and Cold in Arabidopsis. iScience 23.

Garcia-Molina, A. and Pastor, V. (2024). Systemic analysis of metabolome reconfiguration in Arabidopsis after abiotic stressors uncovers metabolites that modulate defense against pathogens. Plant Commun. 5: 100645.

Grant, C.E., Bailey, T.L., and Noble, W.S. (2011). FIMO: scanning for occurrences of a given motif. Bioinformatics 27: 1017–1018.

Hadj-Amor, K., Léger, O., Raffaele, S., Kovacic, M., Duclos, A., Carvalho-Silva, P., Garcia, F., and Barbacci, A. (2024). “Out of sight, out of mind”. The antagonistic role of the transcriptional memory associated with repeated acoustic stimuli on the priming of plant defense. bioRxiv: 2024.05.03.592392.

Hepworth, J. et al. (2020). Natural variation in autumn expression is the major adaptive determinant distinguishing Arabidopsis FLC haplotypes. Elife 9: e57671.

Huot, B., Castroverde, C.D.M., Velásquez, A.C., Hubbard, E., Pulman, J.A., Yao, J., Childs, K.L., Tsuda, K., Montgomery, B.L., and He, S.Y. (2017). Dual impact of elevated temperature on plant defence and bacterial virulence in Arabidopsis. Nat. Commun. 8: 1–11.

Jallet, A.J., Le Rouzic, A., and Genissel, A. (2020). Evolution and Plasticity of the Transcriptome Under Temperature Fluctuations in the Fungal Plant Pathogen Zymoseptoria tritici. Front. Microbiol. 11.

Kappel, C., Friedrich, T., Oberkofler, V., Jiang, L., Crawford, T., Lenhard, M., and Bäurle, I. (2023). Genomic and epigenomic determinants of heat stress-induced transcriptional memory in Arabidopsis. Genome Biol. 24: 1–23.

Katz, E., Li, J.J., Jaegle, B., Ashkenazy, H., Abrahams, S.R., Bagaza, C., Holden, S., Pires, C.J., Angelovici, R., and Kliebenstein, D.J. (2021). Genetic variation, environment and demography intersect to shape Arabidopsis defense metabolite variation across Europe. Elife 10: 1–25.

Kim, J.H. et al. (2022). Increasing the resilience of plant immunity to a warming climate. Nature 607: 339–344.

Kleine, T. et al. (2021). Acclimation in plants – the Green Hub consortium. Plant J. 106: 23–40.

Van Kleunen, M. and Fischer, M. (2005). Constraints on the evolution of adaptive phenotypic plasticity in plants. New Phytol. 166: 49–60.

Lane, D., Denton-Giles, M., Derbyshire, M., and Kamphuis, L.G. (2019). Abiotic conditions governing the myceliogenic germination of Sclerotinia sclerotiorum allowing the basal infection of Brassica napus. Australas. Plant Pathol. 48: 85–91.

Léger, O., Garcia, F., Khafif, M., Carrere, S., Leblanc-Fournier, N., Duclos, A., Tournat, V., Badel, E., Didelon, M., Le Ru, A., Raffaele, S., and Barbacci, A. (2022). Pathogen-derived mechanical cues potentiate the spatio-temporal implementation of plant defense. BMC Biol. 20: 292.

Levis, N.A. and Pfennig, D.W. (2016). Evaluating ‘Plasticity-First’ Evolution in Nature: Key Criteria and Empirical Approaches. Trends Ecol. Evol. 31: 563–574.

Liu, H., Able, A.J., and Able, J.A. (2022). Priming crops for the future: rewiring stress memory. Trends Plant Sci. 27: 699–716.

Liu, Y., Dang, P., Liu, L., and He, C. (2019). Cold acclimation by the CBF–COR pathway in a changing climate: Lessons from Arabidopsis thaliana. Plant Cell Rep. 38: 511–519.

Love, M.I., Huber, W., and Anders, S. (2014). Moderated estimation of fold change and dispersion for RNA-seq data with DESeq2. Genome Biol. 15: 550.

Matsubara, S. (2018). Growing plants in fluctuating environments: why bother? J. Exp. Bot. 69: 4651–4654.

Mbengue, M., Navaud, O., Peyraud, R., Barascud, M., Badet, T., Vincent, R., Barbacci, A., and Raffaele, S. (2016). Emerging Trends in Molecular Interactions between Plants and the Broad Host Range Fungal Pathogens Botrytis cinerea and Sclerotinia sclerotiorum. Front. Plant Sci. 7: 422.

Mee, J.A. and Yeaman, S. (2019). Unpacking Conditional Neutrality: Genomic Signatures of Selection on Conditionally Beneficial and Conditionally Deleterious Mutations. Am. Nat. 194: 529–540.

Mehrabi, Z., Pironon, S., Kantar, M., Ramankutty, N., and Rieseberg, L. (2019). Shifts in the abiotic and biotic environment of cultivated sunflower under future climate change. OCL - Oilseeds fats, Crop. Lipids 26.

Monroe, J.G., Arciniegas, J.P., Moreno, J.L., Sánchez, F., Sierra, S., Valdes, S., Torkamaneh, D., and Chavarriaga, P. (2020). The lowest hanging fruit: Beneficial gene knockouts in past, present, and future crop evolution. Curr. Plant Biol. 24: 100185.

Navaud, O., Barbacci, A., Taylor, A., Clarkson, J.P., and Raffaele, S. (2018). Shifts in diversification rates and host jump frequencies shaped the diversity of host range among *Sclerotiniaceae* fungal plant pathogens. Mol. Ecol.

Newman, R. and Noy, I. (2023). The global costs of extreme weather that are attributable to climate change. Nat. Commun. 14.

Nicotra, A.B., Atkin, O.K., Bonser, S.P., Davidson, A.M., Finnegan, E.J., Mathesius, U., Poot, P., Purugganan, M.D., Richards, C.L., Valladares, F., and van Kleunen, M. (2010). Plant phenotypic plasticity in a changing climate. Trends Plant Sci. 15: 684–692.

Nishad, A. and Nandi, A.K. (2021). Recent advances in plant thermomemory. Plant Cell Rep. 40: 19–27.

Nuruzzaman, M., Sharoni, A.M., and Kikuchi, S. (2013). Roles of NAC transcription factors in the regulation of biotic and abiotic stress responses in plants. Front. Microbiol. 4: 248.

O’Malley, R.C., Huang, S.C., Song, L., Lewsey, M.G., Bartlett, A., Nery, J.R., Galli, M., Gallavotti, A., and Ecker, J.R. (2016). Cistrome and Epicistrome Features Shape the Regulatory DNA Landscape. Cell 165: 1280–1292.

Olate, E., Jiménez-Gómez, J.M., Holuigue, L., and Salinas, J. (2018). NPR1 mediates a novel regulatory pathway in cold acclimation by interacting with HSFA1 factors. Nat. Plants 4: 811–823.

Olson, M. V (1999). When Less Is More: Gene Loss as an Engine of Evolutionary Change. Am. J. Hum. Genet. 64: 18–23.

Ooka, H. et al. (2003). Comprehensive Analysis of NAC Family Genes in Oryza sativa and Arabidopsis thaliana. DNA Res. 10: 239–247.

Peel, M.C., Finlayson, B.L., and McMahon, T.A. (2007). Updated world map of the Köppen-Geiger climate classification. Hydrol. Earth Syst. Sci. 11: 1633–1644.

Peltier, A.J., Bradley, C.A., Chilvers, M.I., Malvick, D.K., Mueller, D.S., Wise, K.A., and Esker, P.D. (2012). Biology, Yield loss and Control of Sclerotinia Stem Rot of Soybean. J. Integr. Pest Manag. 3: 1–7.

Perchepied, L., Balagué, C., Riou, C., Claudel-Renard, C., Rivière, N., Grezes-Besset, B., and Roby, D. (2010). Nitric oxide participates in the complex interplay of defense-related signaling pathways controlling disease resistance to Sclerotinia sclerotiorum in Arabidopsis thaliana. Mol. Plant-Microbe Interact. 23: 846–860.

Roux, F., Voisin, D., Badet, T., Balagué, C., Barlet, X., Huard-Chauveau, C., Roby, D., and Raffaele, S. (2014). Resistance to phytopathogens e tutti quanti: Placing plant quantitative disease resistance on the map. Mol. Plant Pathol. 15: 427–432.

Sadhukhan, A., Prasad, S.S., Mitra, J., Siddiqui, N., Sahoo, L., Kobayashi, Y., and Koyama, H. (2022). How do plants remember drought? Planta 256: 1–15.

Saga, H., Ogawa, T., Kai, K., Suzuki, H., Ogata, Y., Sakurai, N., Shibata, D., and Ohta, D. (2012). Identification and Characterization of ANAC042, a Transcription Factor Family Gene Involved in the Regulation of Camalexin Biosynthesis in Arabidopsis. Mol. Plant-Microbe Interact. 25: 684–696.

Sewelam, N., Oshima, Y., Mitsuda, N., and Ohme-Takagi, M. (2014). A step towards understanding plant responses to multiple environmental stresses: a genome-wide study. Plant. Cell Environ. 37: 2024–2035.

Shahnejat-Bushehri, S., Mueller-Roeber, B., and Balazadeh, S. (2012). Arabidopsis NAC transcription factor JUNGBRUNNEN1 affects thermomemory-associated genes and enhances heat stress tolerance in primed and unprimed conditions. Plant Signal. Behav. 7: 1518–1521.

Shahoveisi, F., Riahi Manesh, M., and del Río Mendoza, L.E. (2022). Modeling risk of Sclerotinia sclerotiorum-induced disease development on canola and dry bean using machine learning algorithms. Sci. Rep. 12: 1–10.

Singh, B.K., Delgado-Baquerizo, M., Egidi, E., Guirado, E., Leach, J.E., Liu, H., and Trivedi, P. (2023). Climate change impacts on plant pathogens, food security and paths forward. Nat. Rev. Microbiol.: 1–17.

Sloat, L.L., Davis, S.J., Gerber, J.S., Moore, F.C., Ray, D.K., West, P.C., and Mueller, N.D. (2020). Climate adaptation by crop migration. Nat. Commun. 11: 1–9.

Somssich, M. (2019). A short history of Arabidopsis thaliana (L.) Heynh. Columbia-0. PeerJ Prepr. 7: e26931v5.

Steinberg, B. and Ostermeier, M. (2024). Environmental changes bridge evolutionary valleys. Sci. Adv. 2: e1500921.

Sucher, J., Mbengue, M., Dresen, A., Barascud, M., Didelon, M., Barbacci, A., and Raffaele, S. (2020). Phylotranscriptomics of the pentapetalae reveals frequent regulatory variation in plant local responses to the fungal pathogen sclerotinia sclerotiorum. Plant Cell 32: 1820–1844.

Torkamaneh, D., Laroche, J., Valliyodan, B., O’Donoughue, L., Cober, E., Rajcan, I., Abdelnoor, R.V., Sreedasyam, A., Schmutz, J., Nguyen, H.T., and Belzile, F. (2019). Soybean Haplotype Map (GmHapMap): A Universal Resource for Soybean Translational and Functional Genomics. bioRxiv: 534578.

Uloth, M.B., You, M.P., Cawthray, G., and Barbetti, M.J. (2015). Temperature adaptation in isolates of Sclerotinia sclerotiorum affects their ability to infect Brassica carinata. Plant Pathol. 64: 1140–1148.

Valladares, F. et al. (2014). The effects of phenotypic plasticity and local adaptation on forecasts of species range shifts under climate change. Ecol. Lett. 17: 1351–1364.

Weng, X., Haque, T., Zhang, L., Razzaque, S., Lovell, J.T., Palacio-Mejía, J.D., Duberney, P., Lloyd-Reilley, J., Bonnette, J., and Juenger, T.E. (2022). A Pleiotropic Flowering Time QTL Exhibits Gene-by-Environment Interaction for Fitness in a Perennial Grass. Mol. Biol. Evol. 39: msac203.

Wood, D.P., Holmberg, J.A., Osborne, O.G., Helmstetter, A.J., Dunning, L.T., Ellison, A.R., Smith, R.J., Lighten, J., and Papadopulos, A.S.T. (2023). Genetic assimilation of ancestral plasticity during parallel adaptation to zinc contamination in Silene uniflora. Nat. Ecol. Evol. 7: 414–423.

Wu, A. et al. (2012). JUNGBRUNNEN1, a reactive oxygen species-responsive NAC transcription factor, regulates longevity in Arabidopsis. Plant Cell 24: 482–506.

Xu, Y.-C. and Guo, Y.-L. (2020). Less Is More, Natural Loss-of-Function Mutation Is a Strategy for Adaptation. Plant Commun. 1: 100103.

Xu, Y.-C., Niu, X.-M., Li, X.-X., He, W., Chen, J.-F., Zou, Y.-P., Wu, Q., Zhang, Y.E., Busch, W., and Guo, Y.-L. (2019). Adaptation and Phenotypic Diversification in Arabidopsis through Loss-of-Function Mutations in Protein-Coding Genes. Plant Cell 31: 1012–1025.

Zandalinas, S.I. and Mittler, R. (2022). Plant responses to multifactorial stress combination. New Phytol. 234: 1161–1167.

Zarattini, M., Farjad, M., Launay, A., Cannella, D., Soulié, M.-C., Bernacchia, G., and Fagard, M. (2021). Every cloud has a silver lining: how abiotic stresses affect gene expression in plant-pathogen interactions. J. Exp. Bot. 72: 1020–1033.

Zheng, F., Zhang, S., Churas, C., Pratt, D., Bahar, I., and Ideker, T. (2021). HiDeF: identifying persistent structures in multiscale ‘omics data. Genome Biol. 22.

Zhong, Z. et al. (2023). Reversed asymmetric warming of sub-diurnal temperature over land during recent decades. Nat. Commun. 14: 7189.

Zuo, D.D., Ahammed, G.J., and Guo, D.L. (2023). Plant transcriptional memory and associated mechanism of abiotic stress tolerance. Plant Physiol. Biochem. 201: 107917.

